# Crystal structure of a bacterial photoactivated adenylate cyclase determined at room temperature by serial femtosecond crystallography

**DOI:** 10.1101/2024.04.21.590439

**Authors:** Sofia M. Kapetanaki, Nicolas Coquelle, David von Stetten, Martin Byrdin, Ronald Rios-Santacruz, Richard Bean, Johan Bielecki, Mohamed Boudjelida, Zsuzsana Fekete, Geoffrey W. Grime, Huijong Han, Caitlin Hatton, Sravya Kantamneni, Konstantin Kharitonov, Chan Kim, Marco Kloos, Faisal H.M. Koua, Iñaki de Diego Martinez, Diogo Melo, Lukas Rane, Adam Round, Ekaterina Round, Abhisakh Sarma, Robin Schubert, Joachim Schulz, Marcin Sikorski, Mohammad Vakili, Joana Valerio, Jovana Vitas, Raphael de Wijn, Agnieszka Wrona, Ninon Zala, Arwen Pearson, Katerina Dörner, Giorgio Schirò, Elspeth F. Garman, András Lukács, Martin Weik

## Abstract

OaPAC is a recently discovered <u>b</u>lue-light <u>u</u>sing flavin adenosine dinucleotide (BLUF) photoactivated adenylate cyclase from the cyanobacterium *Oscillatoria acuminata* that uses adenosine triphosphate and translates the light signal into the production of cyclic adenosine monophosphate. Here, we report the crystal structures of the enzyme in the absence of its natural substrate determined from room temperature serial crystallography data collected at both an X-ray free electron laser and a synchrotron and we compare them with the cryo macromolecular crystallography structures obtained at a synchrotron by us and others. These results reveal slight differences in the structure of the enzyme due to data collection at different temperatures and X-ray sources. We further investigate the effect of the Y6 mutation in the blue-light using flavin adenosine dinucleotide domain, a mutation which results in a rearrangement of the hydrogen-bond network around the flavin and a notable rotation of the side-chain of the critical Q48 residue. These studies pave the way for ps - ms time-resolved serial crystallography experiments at X-ray free electron lasers and synchrotrons in order to determine the early structural intermediates and correlate them with the well-studied ps - ms spectroscopic intermediates.

**Synopsis:** Structures of the dark-adapted state of a photoactivated adenylate cyclase are determined from serial crystallography (SX) data collected at room temperature at an X-ray free electron laser (XFEL) and a synchrotron and are compared with cryo macromolecular crystallography (MX) synchrotron structures obtained by us and others. These structures of the wild-type enzyme in combination with the cryo MX synchrotron structure of a light-sensor domain mutant provide insight into the hydrogen bond network rearrangement upon blue-light illumination and pave the way for the determination of structural intermediates of the enzyme by time-resolved SX.

## 1. Introduction

In the search of light-sensing proteins for optogenetic applications (Yawo *et al*., 2021), <u>p</u>hotoactivated <u>a</u>denylate <u>c</u>yclases (PACs) have emerged as very promising candidates in the field, combining the function of a photoreceptor with that of an enzyme capable of a second messenger generation (Iseki & Park, 2021). First discovered in the unicellular flagellate *Euglena gracilis* (Iseki *et al.,* 2002) to cause a change in the flagellar activity via activation of a protein kinase (Häder & Iseki, 2017) and later identified in both prokaryotes and eukaryotes, PACs are flavoproteins which accelerate the conversion of adenosine triphosphate (ATP) to cyclic adenosine monophosphate (cAMP) upon illumination with blue light (Iseki *et al*. 2002; Stierl, *et al.,* 2011; Penzkofer *et al.,* 2015; Raffelberg *et al.,* 2013; Blain-Hartung *et al.,* 2018; Ryu *et al.,* 2014; Ryu *et al*. 2010). cAMP is a key second messenger in numerous signal transduction pathways, regulating various cellular functions including cell growth, differentiation and motility, gene transcription and protein expression (Gancedo, 2013). In cyanobacteria, cAMP that is produced after blue-light illumination has been shown to bind to a complex involved in the biogenesis of pili required for cell motility (Ohmori & Okamoto, 2004; Yoshimura *et al*., 2002). Hence, cyanobacteria actively sense light gradients which guide their locomotion, moving into or out of areas of favorable conditions (Hoiczyk, 2000). Modulating the cellular concentration of cAMP is of great interest in the field of optogenetics (Ryu *et al.,* 2010; Fomicheva *et al.,* 2019; Stüven *et al*., 2019).

OaPAC is a photoactivated adenylate cyclase from the cyanobacterium *Oscillatoria acuminata* (Ohki *et al*., 2016) and it has a high sequence identity (57%) with the photoactivated adenylate cyclase bPAC from the soil bacterium *Beggiatoa* (Lindner *et al*., 2017). OaPAC is a homodimer of a 366-residue protein comprising an N-terminal BLUF domain and a C-terminal class III adenylyl cyclase (AC) domain. BLUF domains contain a flavin adenine dinucleotide (FAD) chromophore and are ubiquitous blue-light sensors found in bacteria and unicellular eukaryotes. Upon blue-light illumination the hydrogen bond network around the flavin is rearranged resulting in a characteristic 10 nm red-shift in the main visible absorption band of the flavin (Park & Tame, 2017) with the signalling state decaying back in seconds to the dark-adapted state (Ohki *et al.,* 2017). This 10 nm red-shift is a feature not observed in other flavin-based photoreceptors (Chaves *et al*., 2011; Losi *et al.,* 2015; Conrad *et al.,* 2014). The BLUF-regulated AC domain in OaPAC stimulates turnover of ATP up to 20-fold upon illumination (Ohki *et al*. 2016; Blain-Hartung *et al.,* 2018). Synchrotron cryo-crystallography structures of OaPAC in the dark-adapted state have been obtained, both in the presence and the absence of a non-hydrolyzable ATP analogue, as well as in the light-adapted state in the presence of the same analogue (Ohki *et al.,* 2016; Ohki *et al.,* 2017). Recently the room-temperature (RT) and cryo structures of the dark-adapted state of OaPAC in complex with ATP have been solved by serial femtosecond crystallography (SFX) at an X-ray free electron laser (XFEL) and by macromolecular crystallography (MX) at a synchrotron, respectively (Chretien *et al.,* 2024). OaPAC has been shown to be catalytically active in the crystal form as indicated by the characteristic 10 nm red-shift upon blue-light illumination and the increased cAMP levels measured on blue-light exposed crystals soaked in buffer containing ATP and magnesium (Ohki *et al*., 2017). Comparison of the light-and dark-adapted states has shown that the photoactivation mechanism *in crystallo* involves only very small movements suggesting that very small but concerted shifts trigger the enzymatic activity tens of Angstroms from the flavin chromophore (Ohki *et al.,* 2017). However, evidence for a M92_out_/W90_in_ switch major movement has been shown in the recently published crystal structure of ATP-bound OaPAC solved from macrocrystals flash-cooled after 5 sec of blue-light illumination at RT (Chretien *et al*., 2024). In addition, structures solved from microsecond time-resolved (TR) SFX data indicate significant changes around the flavin with the rotation of the side-chain of Q48 (180° at 1.8 μs and 220° at 2.3μs) being the most notable. This rotation is accompanied by a hydrogen bond network rearrangement and destabilization of the loop region between the *β*5/*β*4 strands and the *β*5 strand/*α*3 helix (Chretien *et al*., 2024). Changes in the vibrational modes of the protein backbone between the dark- and light-adapted states of OaPAC, in particular in the *β*5 strand of the BLUF domain which is considered important for signal transduction, have been revealed previously by solution steady-state infrared difference spectroscopy, whereas time-resolved infrared measurements have pointed towards secondary structural changes in the AC domain taking place beyond 100 μs (Collado *et al*., 2022). Indeed, a fast and a slow phase with time constants of 2.3 ms and 36 ms, respectively, have been revealed by time-resolved absorption (TA) and transient grating (TG) experiments (Tokonami *et al*., 2022, Nakasone *et al*., 2023). These phases have been attributed to conformational changes in the BLUF domain, facilitated by W90 and an alteration in the relative orientation of the AC domains for functional activation, respectively.

Despite the present availability of several static structures of dark- and light-adapted states (Ohki *et al*., 2016, 2017, Chretien *et al.,* 2024), structural intermediates in the microsecond timescale (Chretien, A. *et al*., 2024) and solution studies (Collado *et al.,* 2022, Zhou *et al.,* 2022, Tokonami *et al*., 2022, Nakasone *et al*., 2023), the signal transduction pathway in OaPAC has not been revealed. For two reasons, time-resolved SFX is expected to play an essential role in filling this knowledge gap. Firstly, it will allow access to the unexplored ps – ns time scale (Brändén & Neutze, 2021; Weik & Domratcheva, 2022; de Wijn *et al*., 2022, Khusainov *et al*., 2024; Caramello & Royant, 2024), and secondly, it will allow for X-ray radiation damage free data collection (Barends *et al*., 2022; Garman & Weik, 2023). For both reasons, flavoenzymes benefit from TR-SFX, as their structures are prone to radiation damage (Kort *et al.,* 2004) with the bending of the isoalloxazine moiety used as an indicator of the reduction of the flavin (Røhr *et al*., 2010). In addition, flavin radical intermediates are too short-lived to be identified at current synchrotron sources (Maestre-Reyna *et al*., 2022; Sorigué *et al*., 2021).

Here we report the first crystal structures of a substrate-free wild-type OaPAC obtained by RT serial crystallography at both an XFEL and a synchrotron. We compare these structures to substrate-free cryo-MX synchrotron structures of the WT and a mutant in which a tyrosine residue (Y6) that is important for photoactivation is replaced by a tryptophan residue. The latter structure shows the importance of the specific residue on the rearrangement of the hydrogen bond network around the flavin.

## 2. Methods

### 2.1. Protein purification

Full-length wild-type (WT) OaPAC and Y6W OaPAC mutant were expressed and purified as described previously (Collado *et al*., 2022). OaPAC WT and OapAC Y6W plasmids were purchased from GenScript and both sequences were optimized using GenSmart. The WT OaPAC/pET15b or OaPAC Y6W/pET15b plasmid was transformed into BL21(DE3) *E. coli* and grown on a Luria Broth (LB) agar plate containing 100 µg/mL ampicillin. A single colony was used to inoculate 10 mL of LB medium containing 100 µg/ml ampicillin and was grown at 37°C for 16 hours/overnight. This culture was used to inoculate 1 L of LB/ampicillin medium in a 4-L flask. The 1-L culture was incubated at 37 °C until the optical density at 600 nm (OD_600_) reached 0.4-0.5, after which the temperature was decreased to 18 °C followed by the addition of 1 mM isopropyl-b-D-1-thiogalactopyranoside (IPTG) to induce protein expression. The cells were harvested after overnight incubation by centrifugation (6000 rcf, 4 °C) and the cell pellet was stored at −20 °C. The cell pellet was then resuspended in lysis buffer containing protease cocktail inhibitor, DNase (1 mg/mL), 5 mM MgCl_2_ and lysozyme (1 mg/mL). The resuspended cells were lysed by sonication and the cell debris was removed by centrifugation (39,000 rcf, 30 min). The supernatant was loaded onto a Ni-NTA (Qiagen) column, which was washed with buffer containing 20 mM imidazole, and the protein was eluted with wash buffer using 300 mM imidazole. The fractions containing protein were pooled together and purified to homogeneity using size-exclusion chromatography (Superdex-200) and the purity was assessed by SDS-PAGE. The protein concentration was determined using the extinction coefficients ε_280nm_ = 28,590 M^-1^cm^-1^ and ε_445nm_ = 11,300 M^-1^cm^-1^. The following buffers were used: lysis/wash buffer (50 mM Tris pH 8.00, 300 mM NaCl, 20 mM imidazole), elution buffer (50 mM Tris pH 8.0, 300 mM NaCl, 300 mM imidazole) and gel-filtration buffer (50 mM Tris pH 8.0, 150mM NaCl).

### 2.2. Crystallization

Crystallization experiments were carried out at the High Throughput Crystallisation Laboratory (HTX Lab) of the EMBL in Grenoble to find alternative crystallization conditions to those reported in the literature (Ohki *et al*., 2016; 2017) to obtain OaPAC crystals (without substrate added) that diffracted to a higher resolution (<2.9 Å resolution, Ohki *et al*., 2016). OaPAC (7 mg/mL) dissolved in 50 mM Tris 8.0, 150 mM NaCl crystallized in 0.06 M divalents, 0.1 M Tris-Bicine pH 8.5, 30% v/v PEG550 MME-PEG20000 at 20°C. Under these conditions, crystals reached dimensions of 90 μm × 50 μm × 20 μm within two days and were prepared for X-ray diffraction experiments using the CrystalDirect technology (Zander *et al*., 2016) and cryo-MX synchrotron data were collected at 100 K. Having identified initial macrocrystallization conditions, OaPAC microcrystals (without substrate added) were produced at 20°C using a batch crystallization approach without seeding. 300 μl OaPAC (7 mg/mL) was mixed with 700 μl crystallization solution (0.06 M divalents, 0.1 M Tris-Bicine pH 8.5, 30% v/v PEG500 MME-PEG20000) and left in the dark. The size of the microcrystals (<16 μm × 15 μm × 5 μm was estimated using a Discovery V12 Stereo Zeiss microscope.

### 2.3 Data collection and processing, structure solution and refinement

SFX experiments were carried out at the SPB/SFX instrument (Mancuso *et al*., 2019) of the European XFEL (Schenefeld, Germany) (proposal PCS 3018, 28-29 October 2022, 4.5 hours). The OaPAC microcrystals jetting conditions were tested in the injection lab at IBS and in the XFEL Biology Infrastructure (XBI) lab (Han *et al*., 2021) before the beamtime. A total volume of 4.5 mL of OaPAC microcrystal slurry (10% v/v gravity-settled crystals) (Figure 1i) was injected into the Interaction Region Upstream (IRU) under high vacuum with a standard gas dynamic virtual nozzle (GDVN) (75 μm inner diameter) (Vakili *et al*., 2022). The GDVN was equipped with a 30 x 34 μm inline sample filter, and operated with liquid flow rate of 30 μl/min (corresponding to a jet velocity of 45 m/s). The sample was probed using trains delivered at 10 Hz with a 564 kHz intra-train repetition rate (9.3 keV photon energy, 50 fs (FWHM) pulse length, ∼2.5 mJ pulse energy), each containing 180 pulses spaced by 1.773 μs in time. The X-ray focal size was 3 μm x 3 μm diameter (FWHM) and the time interval between each 180-pulse train was ∼100 ms. Radiation damage free diffraction data (Nass *et al*., 2019) from OaPAC crystals was recorded using a 1-megapixel Adaptive Gain Integrating Pixel Detector (AGIPD) (Allahgholi *et al.,* 2019). To avoid sample damage by the previous pulses (Stan, *et al*., 2016; Grünbein *et al*., 2021), an optical imaging system with an fs optical laser for jet illumination was used to observe the effects of the interaction with the XFEL pulses, and to make sure that the liquid jet had time to recover between two successive X-ray pulses.

**Figure 1.**
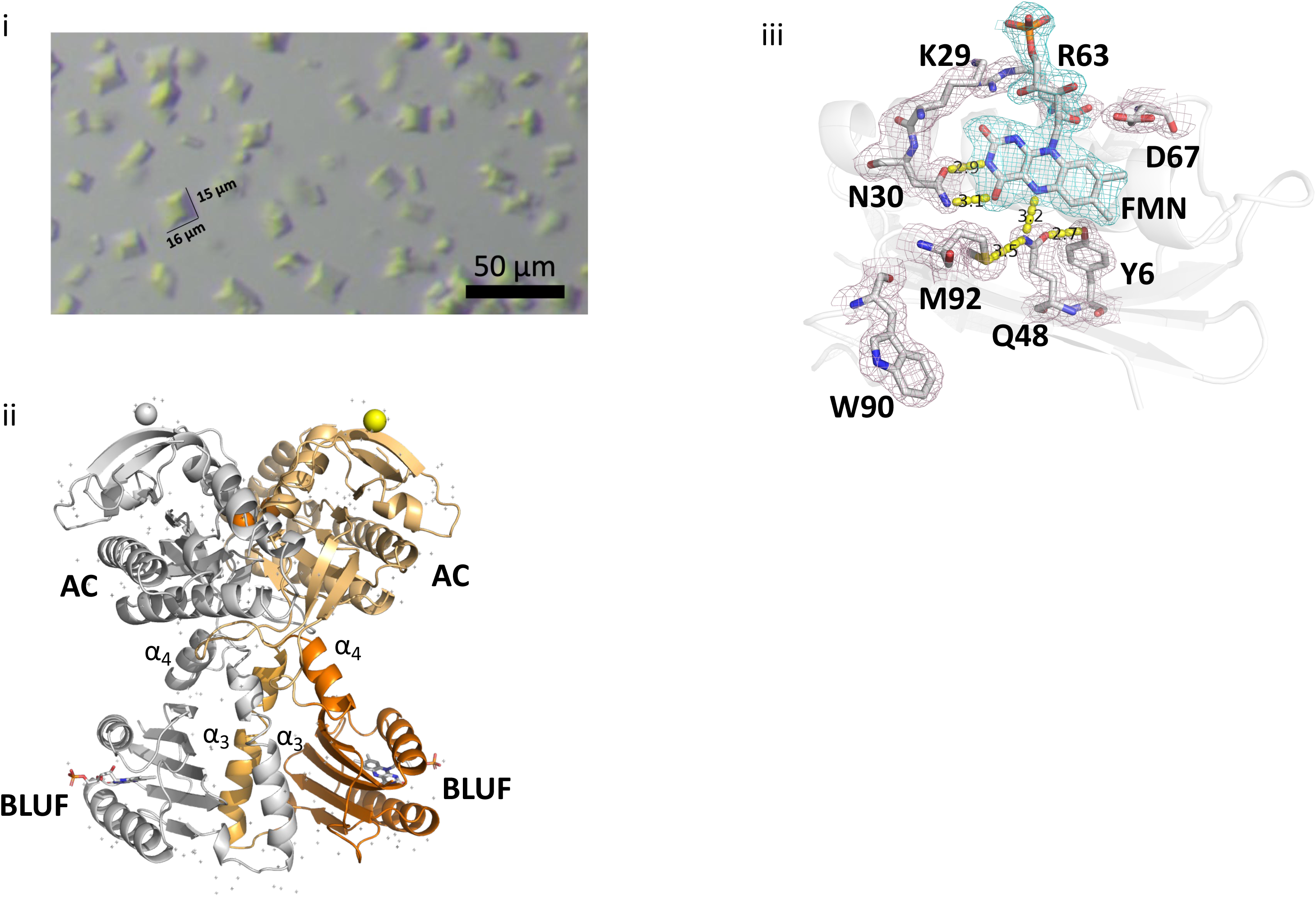
i) OaPAC crystals used at the EuXFEL. Crystal size <16 μm x 15 μm x 5 μm; the scalebar corresponds to 50 μm ii) Ribbon diagram of the SFX structure of wild-type; monomeric OaPAC in grey; BLUF domain, orange; α_3_ helix, light orange; handle: α_4_ helix, bright orange; AC domain, yellow orange; calcium ions are represented by spheres iii) Flavin active site of OaPAC in the SFX structure showing amino acid residues participating in the hydrogen bond network. 2Fo-Fc electron density maps are represented in blue mesh (flavin) and light purple (residues) at a contour level of 1.0 σ.

Hits were identified using NanoPeakCell (Coquelle *et al.,* 2015) and saved into h5 files for further processing (indexing and integration) with CrystFEL v 0.10.1 (White, 2018). A total of 17,434,418 raw images were collected with a hit rate of 1.6 % (number of hits 285,671) consuming 4.5 mL of a microcrystal suspension. Diffraction spots were detected at a resolution better than 1.75 Å. 27,202 lattices were indexed (indexing rate of 9.5 %) in the space group P2_1_2_1_2 with unit-cell parameters a=102.90 Å, b=54.91 Å, c=72.98 Å, α=β=γ=90°, (Table1). By inspection of the R_split_ and CC_1/2_ values the high-resolution limit was determined to be 1.75 Å.

For structure refinement, starting phases were obtained by molecular replacement using *Phenix* (Liebschner *et al*., 2019). The search model was the PDB entry 4yut (Ohki *et al*., 2016) after removing the flavin mononucleotide (FMN; isoalloxazine ring with a ribityl and one phosphate group). It should be noted that although BLUF domains *in vivo* are assumed to contain FAD (Laan *et al*., 2004), electron density corresponding to the adenine moiety of FAD has not been identified in OaPAC expressed in *E. coli* (Ohki *et al.,* 2016) similar to other BLUF domains (AppA, bPAC) (Anderson *et al*., 2005, Jung *et al*., 2006, Lindner *et al*., 2017). Refinement in real and reciprocal space, electron-density map calculations and solvent-molecule assignments were performed using *Phenix*. The occupancies of the alternate conformations found in several residues were refined using occupancy refinement in *Phenix* after initial values were set to equal occupancy (0.5) for each conformer. Model building and real space refinement was performed using *Coot* (Emsley *et al*., 2010). During the refinement process, water molecules, FMN and calcium ions were added stepwise to the model. *PyMOL* (Schrödinger, New York, USA) was used to prepare figures. Data collection and refinement statistics are summarized in Table 1. Atomic coordinates and structure factors have been deposited in the PDB (accession codes 9f1w, 9f1x, 9f1y and 9f1z).

**Table 1.**
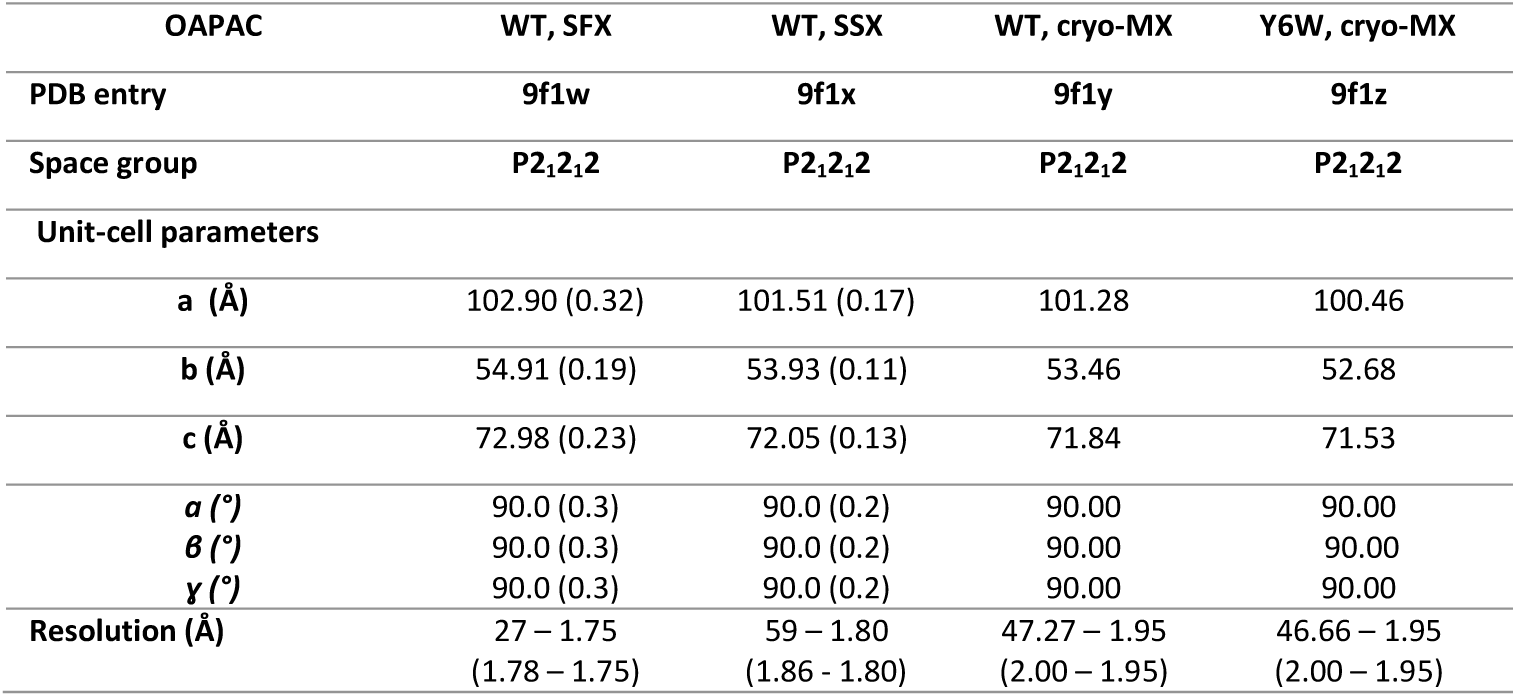
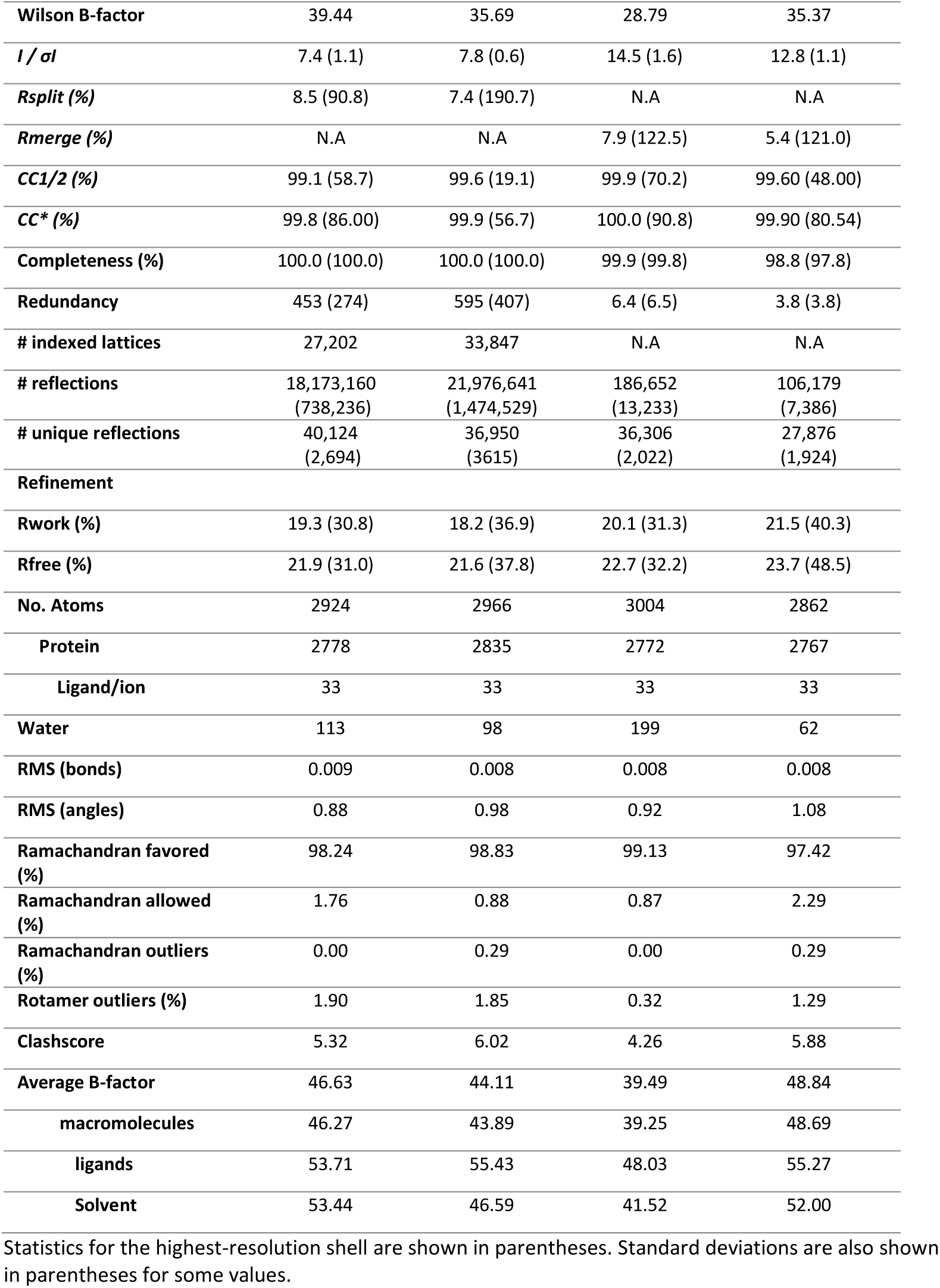
Data collection and refinement statistics for WT-type OaPAC and Y6W OaPAC mutant.

The synchrotron serial X-ray diffraction experiment was conducted at RT at the EMBL beamline P14-2 (T-REXX) at the PETRA III synchrotron (DESY, Hamburg) using a 12.7 keV beam with a flux of ∼2 x 10^12 ph/s and a diameter of 10 μm. The 16 μm x 15 μm x 5 μm microcrystals were loaded onto the HARE chips and mounted on the translation-stage holders as described previously (Mehrabi *et al*., 2020). Diffraction images were processed automatically at the beamline using CrystFEL v 0.10.2 (White, 2018) with the XGANDALF indexing routine (Gevorkov *et al*., 2019). Diffraction data from the wild-type and Y6W OaPAC crystals were collected at 100 K at the ID30A-1 beamline of the ESRF, Grenoble (IBS-BAG mx2398). The SSX structure of wild-type OaPAC was solved by molecular replacement using DIMPLE (CCP4 suite) (Agirre *et al*., 2023) whereas the refinement and the manual rebuilding procedure of the SSX and MX data was similar to that of the SFX one, as described above.

### 2.4 Micro-PIXE

Proton induced X-ray emission (PIXE) analysis of proteins was carried out at the Ion Beam Centre, University of Surrey, UK using the methods detailed in (Grime & Garman, 2023). In summary, characteristic X-ray emission was induced using a 2.5-MeV proton beam of less than 2 µm diameter incident under vacuum on dried OaPAC crystals that had been washed in a metal free buffer (volume per droplet approx. 0.1 μL) before mounting.

The emitted X-rays were detected using a silicon drift detector (Rayspec Ltd, UK). In order to detect sodium and magnesium, we removed the 130 µm thick beryllium filter normally used to avoid deleterious effects of recoiling protons. Additionally, backscattered protons were detected using a silicon surface barrier particle detector and the resulting spectrum was analysed to give the sample thickness and gross matrix composition. By scanning the proton beam in *x* and *y* over the dried sample, spatial maps were obtained of all elements with Z ≥11 (sodium) present in the sample. Quantitative information was obtained by collecting three- or four-point spectra from each droplet. PIXE spectra were processed with GUPIX (Campbell *et al*., 2021) using the matrix composition derived from the simultaneously collected RBS spectrum. The concentrations of the elements were used to calculate the relative amount of each element per protein molecule in the sample using the sulphur signal from the known number of cysteines and methionines in the protein sequence as an internal standard for normalisation.

### 2.5 UV-visible absorption spectroscopy on OaPAC macrocrystals

To examine the photoactivity of OaPAC crystals in the specific crystallization condition used here, we recorded the absorption spectra of OaPAC crystals in the dark- and light-adapted state (Figure S1i) using the Cal(ai)^2^doscope, a custom-made microspectrophotometer (Rane *et al*., 2023). An OaPAC crystal (90 μm × 50 μm × 20 μm) was mounted between two microscope slides and absorption spectra were recorded at RT in the dark-adapted state and in the light-adapted state after illumination with a cw 488 nm laser (illumination for 5 sec, 1 mW power, top hat FWHM 100 μm laser spot). Absorption spectra were also recorded for OaPAC in solution with a SPECORD S600 (AnalytikJena) spectrophotometer (Figure S1ii).

## 3. Results and Discussion

### 3.1. Room temperature SFX structure of substrate-free OaPAC

For this work, the SFX structure of substrate-free OaPAC was solved at 1.75 Å resolution (Figure 1ii). OaPAC crystals (Fig. 1i) were grown in the dark in a crystal form (P2_1_2_1_2; a= 102.90 Å, b=54.91 Å, c=72.98 Å, α=β=γ=90°) with one monomer in the asymmetric unit. Figure 1ii shows a ribbon model with the characteristic parallel dumbbell-shaped dimer with an antiparallel arrangement of the N-terminal BLUF and C-terminal AC domains, as previously described for the cryo-structures (Ohki *et al*., 2016; Ohki *et al*., 2017, Chretien *et al*., 2024) and for the SFX structure of ATP-bound OaPAC (Chretien *et al*., 2024). The BLUF domains dimerize through their α3_BLUF_-capping helices that form an intermolecular coiled coil whereas a four-helix bundle consisting of the C termini of the α3-capping helices, the α4_BLUF_ helix and the α4_BLUF_-β1_AC_ loop is formed on which the AC domains reside. The central coiled-coil domain has been suggested to play a key role in the transduction of the signal received by the BLUF domain (Ohki *et al*., 2016).

In earlier studies, OaPAC was crystallized in orthorhombic (P2_1_2_1_2_1_, a= 85.3 Å, b=100.7 Å, c=120.7 Å, α=β=γ=90° at 100 K, resolution 2.9 Å, pdb 4yut) and hexagonal (P6_1_22, a= 76.8 Å, b=76.8 Å, c=204.8 Å, α=β=90°, γ=120° at 100 K, resolution 1.8 Å, pdb 4yus,) crystal forms (Ohki *et al*., 2016; Ohki *et al.,* 2017). In the latter, the best resolution was achieved in the presence of a nonhydrolyzable ATP analogue (ApCpp) although no electron density was observed for it. Yet, another space group has been recently reported for the cryo-MX structure of OaPAC in the absence of substrate (C222_1_, a=52.8 Å, b=146.2 Å, c=103.6 Å, α=β=γ=90° at 100 K, resolution 1.5 Å, pdb 8qfe), for the cryo-MX structure of the ATP-bound OaPAC (C222_1_, a=54.5 Å, b=146.4 Å, c=104.9 Å, α=β=γ=90° at 100 K, resolution 2.1 Å, pdb 8qff), and the SFX structure of the ATP-bound OaPAC (C222_1_, a=54.3 Å, b=145.8 Å, c=105.3 Å, α=β=γ=90° at RT, resolution 1.8 Å, pdb 8qfh). We attribute the differences in the space group and unit cells to the different crystallization conditions used. In addition, subtle differences are observed in the percentages of the secondary structure elements in our SFX substrate-free OaPAC model: using YASARA (www.yasara.org), we have calculated the following contributions: 42% helix, 29.6% sheet, 7.5% turn, 20.9% coil for it. These values are slightly different to those calculated for the cryo-structures (cryo-, 42.9% helix, 29.1% sheet, 8.9% turn, 19.1% coil; *pdb:4yut*, 41% helix, 26.1% sheet, 7.7% turn, 25.2% coil; *pdb:4yus*, 44.6% helix, 30.6% sheet, 6.6% turn, 18.3% coil; *pdb: 8qfe*, 39.7% helix, 28% sheet, 10.3% turn, 22% coil) and are all summarized in Table 2.

**Table 2.** Contribution of secondary structure elements in all OaPAC structures.

|  | helix (%) | sheet (%) | turn (%) | coil (%) |
| --- | --- | --- | --- | --- |
| **SFX (9f1w)** | 42.0 | 29.6 | 7.5 | 20.9 |
| **SSX (9f1x)** | 41.9 | 29.5 | 7.5 | 21.1 |
| **WT-cryo (9f1y)** | 42.9 | 29.1 | 8.9 | 19.1 |
| **Y6W-cryo (9f1z)** | 42.5 | 29.1 | 7.7 | 20.8 |
| **4yut** | 41.0 | 26.1 | 7.7 | 25.2 |
| **4yus** | 44.6 | 30.6 | 6.6 | 18.3 |
| **Cryo (8qfe)** | 39.7 | 28.0 | 10.3 | 22.0 |
| **Cryo ATP bound (8qff)** | 40.3 | 28.0 | 8.0 | 22.3 |
| **SFX ATP bound (8qfh)** | 40.3 | 28.0 | 8.0 | 23.7 |

Upon blue light illumination, crystals of OaPAC have been previously shown to produce a characteristic red shift and to turn over ATP (Ohki *et al.,* 2017). Using a custom-made microspectrophotometer (Byrdin & Bourgeois, 2016), we examined the photoactivity of the OaPAC crystals produced with our specific crystallization conditions. Figure S1i shows the absorption spectra of an OaPAC crystal in the dark- and in the light-adapted states. The light-adapted state is characterized by the typical red shift observed for OaPAC in solution (FigS1ii). The absorption spectrum of oxidized flavin of OaPAC in solution exhibits two broad absorption bands, with peaks at 444 nm for S_0_=>S_1_ and at 375 nm for S_0_=>S_2_ (Stanley & MacFarlane, 2000) which red shift upon blue blue-light excitation as observed in BLUF domains. The similar red shift observed also in our OaPAC crystals (Figure S1i) suggests that the specific crystallization condition produces crystals that are photoactive and hence suitable for time-resolved crystallographic experiments. Such studies will allow a correlation of the available spectroscopic signatures on the substrate-free enzyme (Collado *et al*., 2022, Tolentino Collado *et al*., 2024) with the early-formed structural intermediates.

### 3.2. Structural analysis of the BLUF domain of the SFX structure of substrate-free OaPAC

The overall fold of the BLUF domain in the RT SFX crystal structure of substrate-free OaPAC reveals the characteristic compact arrangement of BLUF domains. It consists of the five-stranded mixed β-sheet with two α-helices running parallel to the β-strands on one side and the helix-turn-helix unit on the other side of the sheet. The first 96 amino acid residues of the protein form an αβ sandwich with a typical ferredoxin-like β_1_α_1_β_2_β_3_α_2_β_4_ topology (Figure 1ii). Figure 1iii shows the H-bonding interactions in the flavin binding pocket and a clear electron density for the flavin and the ribityl side chain but no apparent density for the adenine moiety, similarly to previous observations in OaPAC (Ohki *et al*., 2016) and other BLUF domains (AppA, bPAC) (Anderson *et al*., 2005, Jung *et al.,* 2006, Lindner *et al*., 2017). In the flavin binding environment we identify the conserved residues Y6 and Q48 that play an important role in the photoactivation mechanism (Collado *et al*., 2022). The amino group of Q48 forms H-bonds with methionine M92 (3.5 Å) and N5 of the flavin (3.2 Å), whereas the carbonyl group forms a H-bond with the hydroxyl group of Y6 (2.7 Å). Other residues that are part of the H-bond network in the active site are i) N30 which H-bonds with the C4=O carbonyl and the N3 of the FMN, ii) R63 which *via* the amino group H-bonds with the C2=O carbonyl of the FMN and via the backbone carbonyl with the O atom of C2’ of the ribityl chain of the FMN (not shown), iii) K29 which H-bonds with one O atom of the phosphate group of the FMN and the O atom of the C4’ of the ribityl chain of the FMN (not shown), and iv) D67 which H-bonds with the O atoms of C2’ and C3’ of the ribityl chain of the FMN (not shown).

### 3.3 Structural analysis of the AC domain of the SFX structure of the substrate-free OaPAC

The AC domains in OaPAC exhibit the canonical type III fold and form a homodimer with two active sites at the dimer interface (Figure 2i). In class III AC domains, the catalytic centers are formed at the dimer interface employing a two-metal ion mechanism where one metal ion coordinates to the triphosphate moiety of ATP while the other ion activates the 3-hydroxy group for nucleophilic attack on the *α*-phosphate (Linder, 2006). Figure 2ii shows two areas at the dimer interface where electron density (not shown) exists between D156/D200/E279 (chain A) and D270/N273 (chain B) and between D156/D200/E279 (chain B) and D270/N273 (chain A). Based on the ATP binding site arrangement in Y7F bPAC (Lindner *et al*., 2017) and in class III AC domains (Steegborn, 2014) this electron density (not shown) can be attributed to the two metal ion binding sites. Indeed, the recently published ATP-bound structure of OaPAC confirms the involvement of the residues D156, D200 and E279 in the ATP binding site that requires two magnesium ions when ATP is bound and one in the ATP-free form (Chretien *et al.,* 2024). Besides these two metal ion binding sites in our SFX structure of the substrate-free OaPAC, we also observe electron density (not shown) around D321/D310 for both chains A and B at the surface of the AC domains (Figure 2iii, iv).

**Figure 2.**
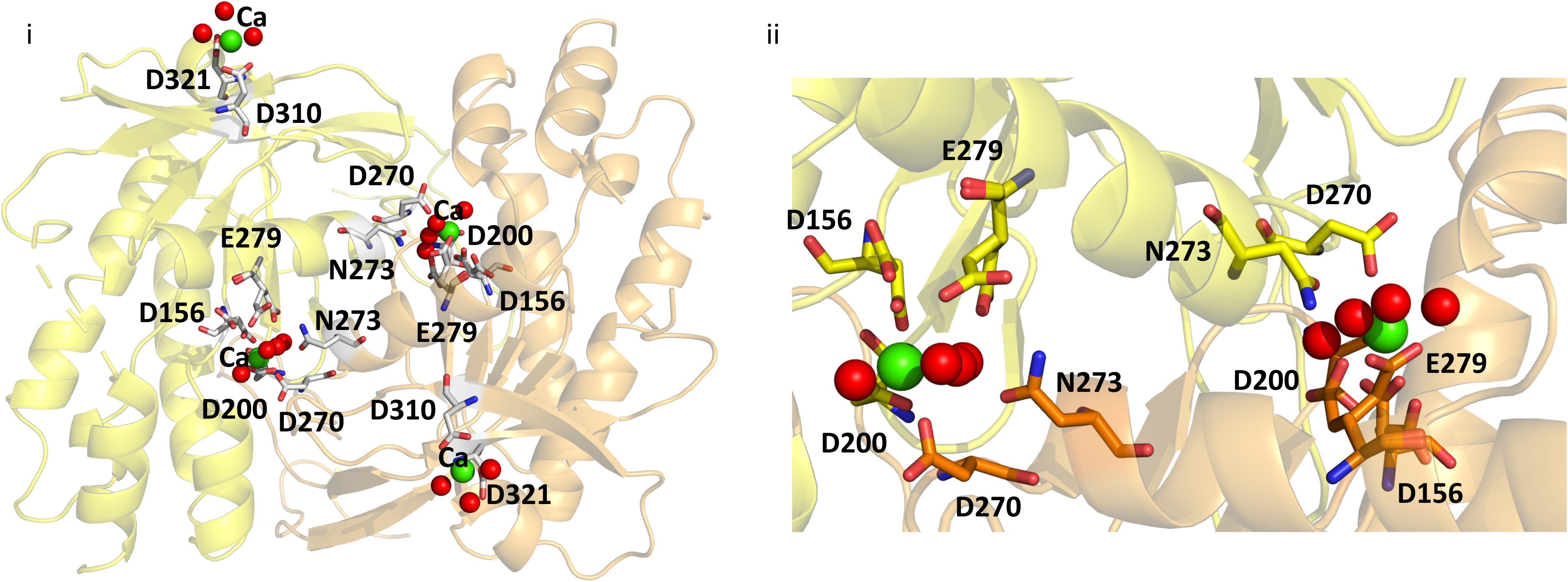
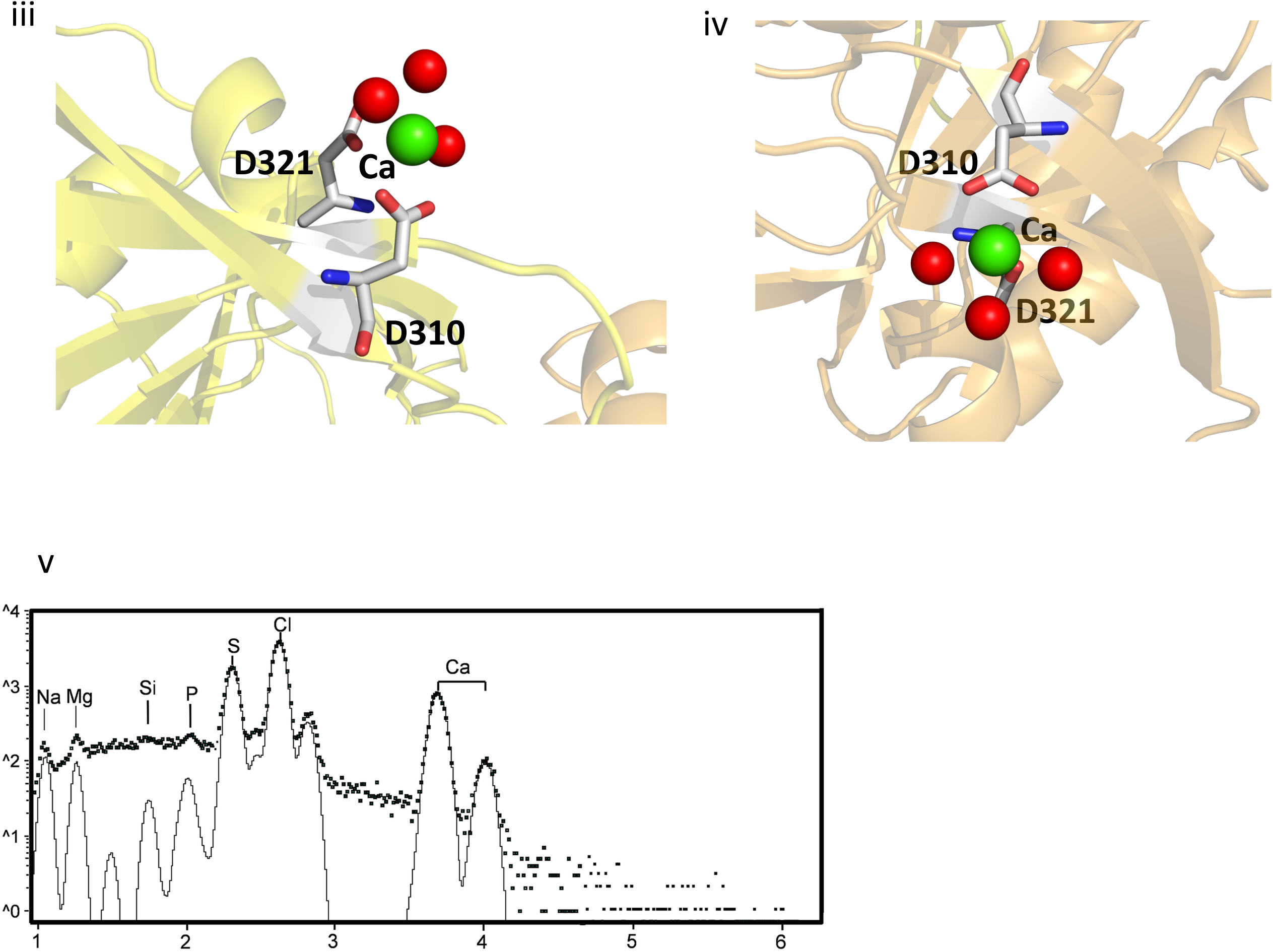
i) AC domains in OaPAC (SFX) showing the metal ion binding sites and surrounding residues; green spheres represent calcium ions and red spheres represent water molecules ii) Zoom-in on the dimer interface and the internal calcium binding sites with coordinating residues; iii) and iv) Zoom-in on the calcium binding sites on the surface of OaPAC v) Micro-PIXE analysis of OaPAC confirms the presence of calcium and magnesium ions in the crystals.

Although our crystallization condition contains both magnesium (Mg) and calcium (Ca) our experimental results (PIXE experiments) and theoretical calculations (CheckMyMetal; https://cmm.minorlab.org) as discussed below suggest that only calcium is present in the binding sites of the AC domains. In particular, Figure 2v shows the micro-PIXE analysis of WT OaPAC crystals which have an average of two Mg atoms and 11 Ca atoms per protein molecule. The signal for the point spectra for Mg was between 3× and 6× above the limit of detection (LOD) and for Ca was between 50× and 100× above the LOD. The *y*-axis is displayed on a log_10_ scale to help to visualize trace elements. Characteristic peaks for sulphur (S), chlorine (Cl) and calcium (Ca) atoms are highlighted. The black line shows the fit to these peaks following background subtraction. In addition, the validation server ‘check my metal’ (Gucwa *et al*., 2022) favors the presence of a calcium atom in each one of the four binding sites in the biological dimer, whereas modeling of magnesium results in large mFo-DFc peaks (Fig. S2) suggesting that ions with more electrons are actually present. Based on the above analysis, we propose that the Mg ions identified in the micro-PIXE analysis may be present in channels of the enzyme. Although Mg ions have been observed in the ATP binding site of the recently reported SFX structure of ATP-bound OaPAC (Chretien *et al.,* 2024) it should be pointed out that there were no Ca ions present in their crystallization condition. Calcium is a non-transition element which demonstrates a variety of coordination numbers with six to eight being the most usual ones and as a ‘hard’ metal ion (Ca^2+^) prefers ‘hard’ ligands with low polarizability, with oxygen being the most preferable coordinating atom (Dudev & Lim, 2003). The identified Ca^2+^ ions coordinate with four water molecules and five residues (D156, D200, D270, N273, E279) in the internal metal binding site whereas on the surface they coordinate with three water molecules and two residues (D310, D321) (Figure 2ii, iii, iv).

### 3.4 Comparison of the SFX structure of substrate-free OaPAC with the SSX and cryo synchrotron structure of substrate-free OaPAC

Our RT SFX structure displays a similar fold as our structures obtained at room temperature (0.496 Å Cα RMSD over 350 residues) and at 100 K (0.508 Å Cα RMSD over 350 residues) at a synchrotron. They also share the same space group (P2_1_2_1_2, Table 1) whereas there are no significant changes in the contribution of the secondary structure elements (Table 2). Similarly, calculation of the theoretical SAXS parameters derived from the crystal structures using the FoXS server (Schneidman-Duhovny *et al*., 2016) also does not reveal significant differences in the radius of gyration R_g_ and D_max_ which are in line with the recently reported values in solution (Ujfalusi-Pozsonyi, *et al.,* 2024) (Table 3, Figure S3). In order to reveal any potential differences between these structures, we have calculated the distance difference matrices (DDM) which highlight changes in interatomic distances. Red and blue colors indicate an increase or decrease in distances, respectively with respect to the reference structure (Seno & Go, 1990). The DDM reveal that the cryo- and the SSX structures are more compact than the SFX structure (Figure 3i, ii). A comparison of the BLUF and AC domains of the SFX OaPAC structure with those of the cryo- and SSX ones are shown in Figure 3iii/iv and Figure 3v/vi, respectively. The SFX OaPAC structure is almost identical to the SSX OaPAC structure as there are hardly any differences observed (Figure 3v/vi). Small differences are observed in the internal cavities of the SFX OaPAC structure compared to the SSX OaPAC and the cryo-OaPAC structure, as illustrated in Figure 3vii, 3viii.

**Figure 3.**
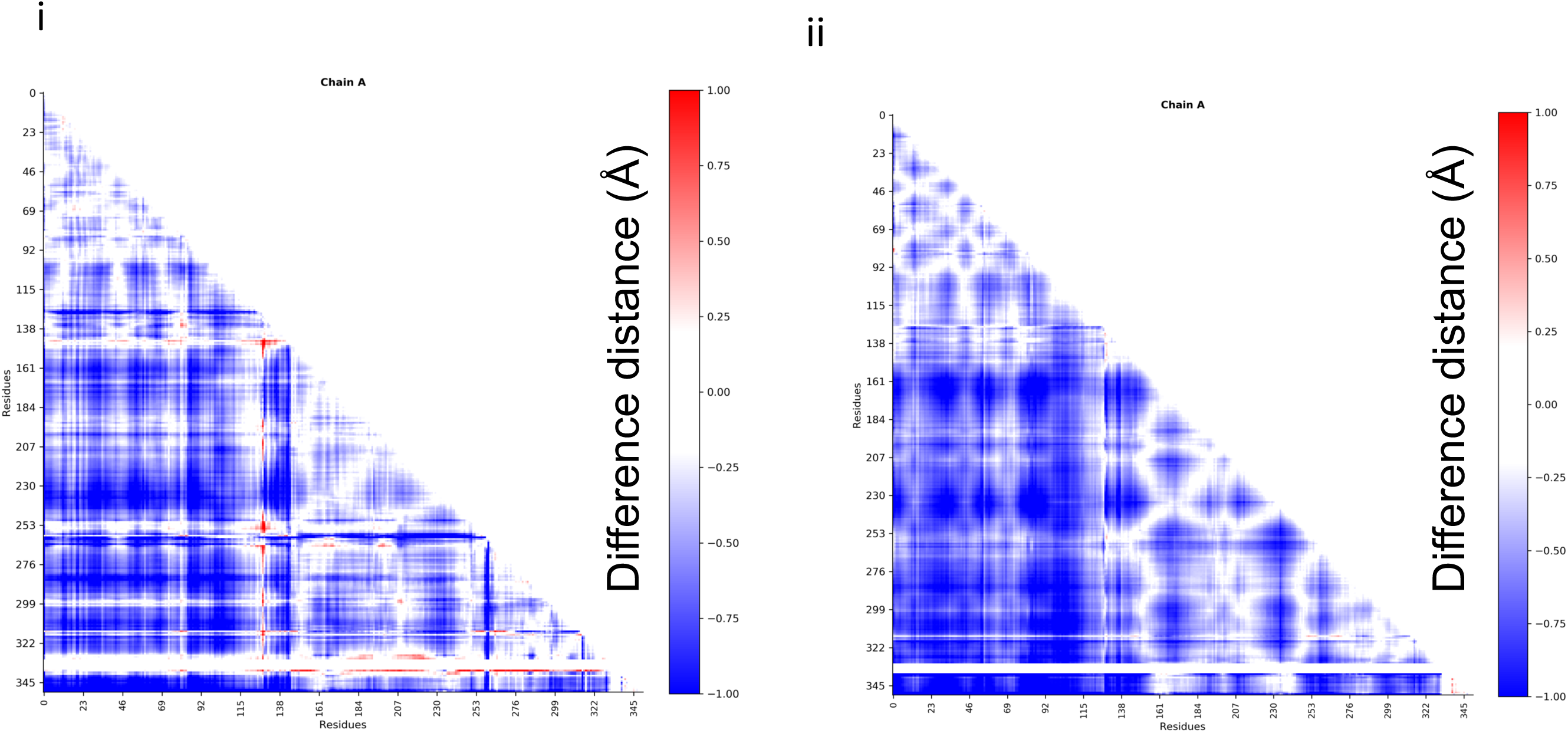
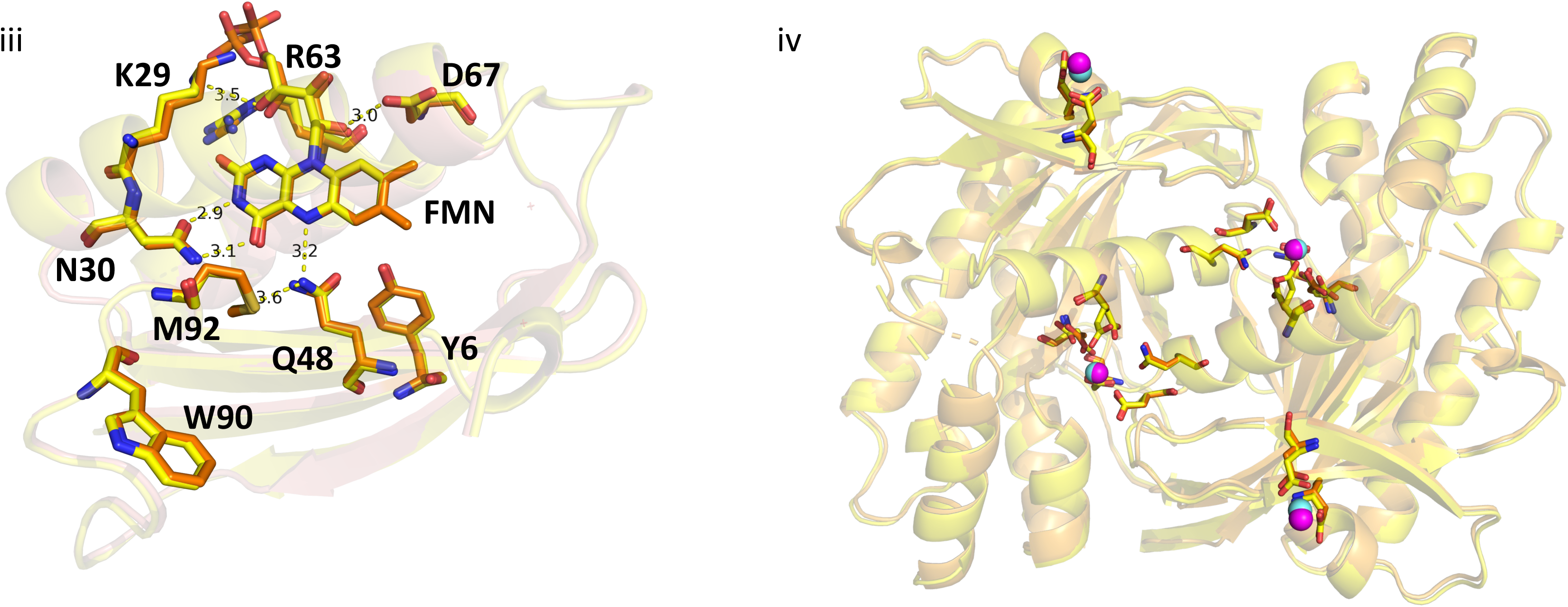
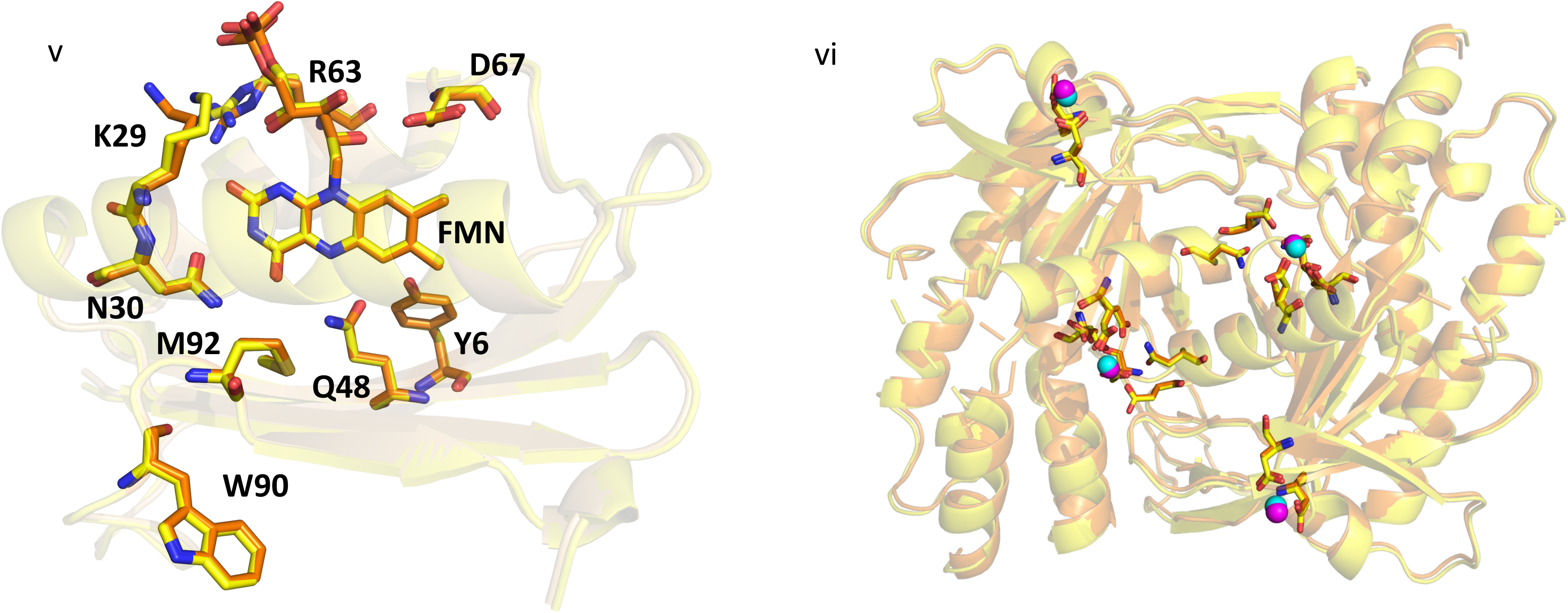
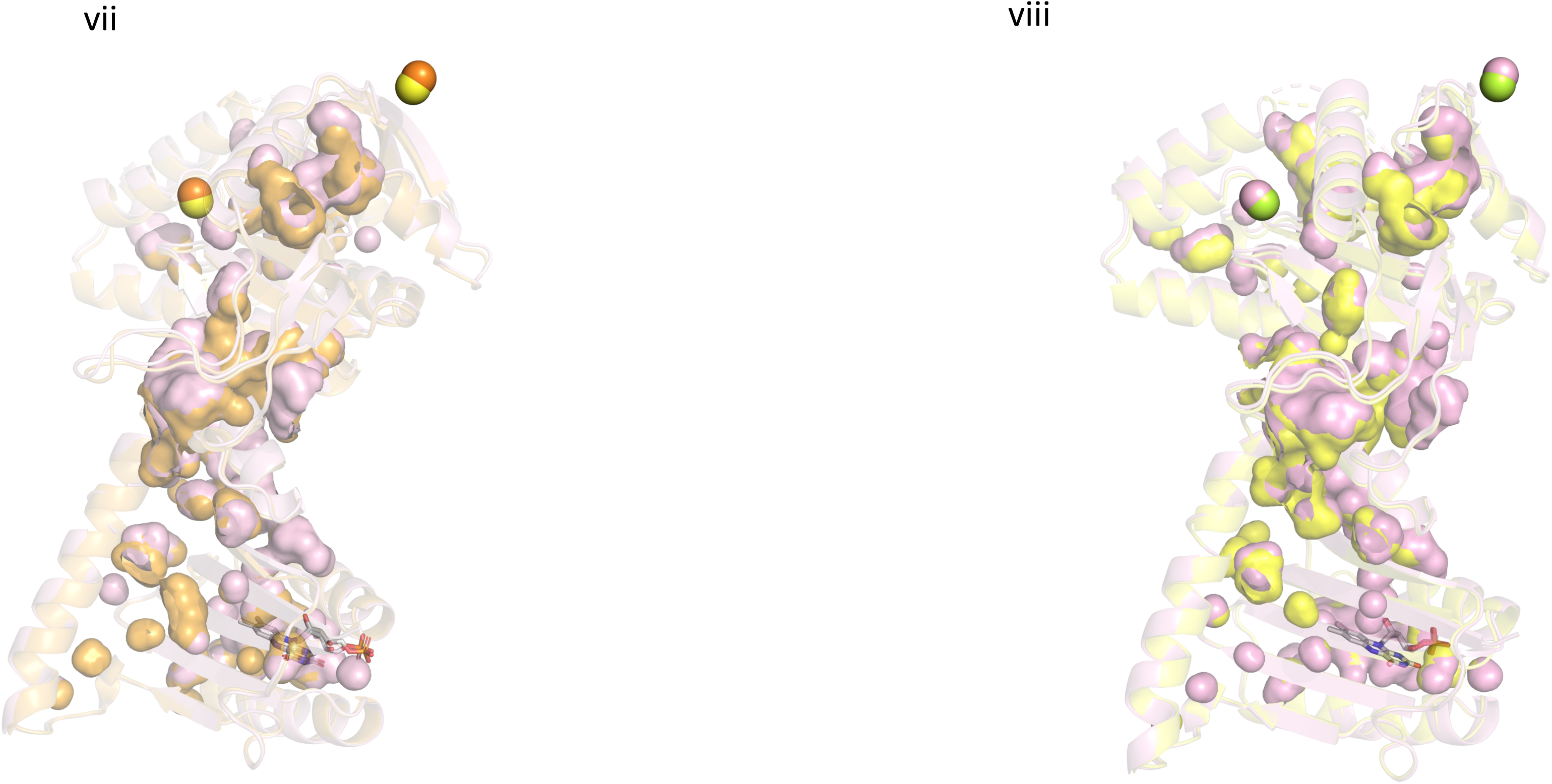
i) Difference distance matrix calculated between the SFX structure and the cryo-structure of OaPAC. Red and blue indicate increasing and decreasing distances, respectively with respect to the SFX structure ii) Difference distance matrix calculated between the SFX structure and the SSX structure of OaPAC. Red and blue indicate increasing and decreasing distances, respectively with respect to the SFX structure. Residues 332-337 are not resolved in the SFX structure which results in the white area in this region in the DDM map iii) Superimposition of the BLUF domains of the SFX structure of OaPAC (yellow) and of the cryo-structure of OaPAC (orange). Distances are shown for the SFX structure of OaPAC iv) Superimposition of the AC domains of the SFX structure of OaPAC (yellow) and of the cryo structure of OaPAC (orange) showing amino acid residues coordinated to the calcium ions (SFX; pink spheres, cryo; cyan spheres) v) Superimposition of the BLUF domains of the SFX structure of OaPAC (yellow) and of the SSX structure of OaPAC (orange) vi) Superimposition of the AC domains of the SFX structure of OaPAC (yellow) and of the SSX structure of OaPAC (orange) showing amino acid residues coordinated to the calcium ions (SFX; pink spheres, cryo; cyan spheres) vii) Superimposition of monomeric of OaPAC (SFX pink; cryo; orange; calcium ions in SFX; orange spheres, calcium ions in cryo; yellow) showing cavities calculated with PyMOL viii) Superimposition of monomeric of OaPAC (SFX pink; SSX; yellow; calcium ions in SFX; pink spheres, calcium ions in SSX; green) showing cavities calculated with PyMOL.

**Table 3.** Basic SAXS parameters R_g_ (radius of gyration), D_max_ (maximum size) and R_g_(GNOM) (radius of gyration calculated using GNOM (Svergun, 1992) derived from the theoretical SAXS profiles of the OaPAC structures (Figure S3).

|  | **SFX** | **SSX** | **cryo** | **Y6W** | **SEC-SAXS\*** |
| --- | --- | --- | --- | --- | --- |
| $R_g(\text{\AA})/r^2$ | 29.0648 ± 0.6438 /1.0 | 28.6412 ± 0.5649 /1.0 | 28.7891 ± 0.5879 /1.0 | 28.5921 ± 0.5723 /1.0 | 31.4 ± 0.52 /0.9908 |
| $R_g(\text{\AA})$ (GNOM) | 29.06 ± 0.0590 | 28.63 ± 0.0512 | 28.78 ± 0.0494 | 28.59 ± 0.0569 | 30.71 ± 0.01 |
| $D_{max}(\text{\AA})$ /$\chi^2$ | 86/0.9185 | 85/0.9191 | 85/0.9183 | 85/0.9195 | 90/1.0728 |

\* Ujfalusi-Pozsonyi, K. *et al.*, 2024.

### 3.5 Comparison of the WT OaPAC and Y6W mutant structures obtained by cryo-MX

Tyrosine 6 plays an important role in the photoactivation mechanism of OaPAC (Collado *et al*. 2022) and of BLUF domains in general (Park & Tame, 2017). In OaPAC, a concerted proton-coupled electron transfer (PCET) from Y6 to form the neutral flavin radical and subsequent recombination results in the hydrogen bond rearrangement around the flavin that eventually causes transduction of the light signal to the AC domains. Mutation of the analogous tyrosine residue in AppA (Y21W), the BLUF containing Activation of Photopigment and pucA photoreceptor from *Rhodobacter sphaeroides* results in the formation of the anion flavin radical compared to the neutral flavin radical formed in the WT (Lukacs *et al*., 2014). In PixD, the BLUF protein from *Synechocystis*, the equivalent mutation (Y8W) blocks the formation of the signaling state (Bonetti *et al*., 2009), whereas in OaPAC_BLUF_, the Y6W mutation was recently introduced to dissect the forward electron and proton transfer following blue-light excitation of the flavin (Chen *et al*., 2023). To investigate the structural role of this residue, we have mutated Y6 to a tryptophan (Y6W), a mutation which is expected to disturb the H-bond network (Figure 1iii) around the flavin and at the same time forms a photoinactive enzyme (Lukacs *et al*. unpublished results). The Y6W mutation does not result in a change of the space group (still P2_1_2_1_2) and unit cell parameters (Table 1) but seems to induce the anticipated change in the H-bond network around the flavin. In particular, loss of the H-bond between Y6 and Q48 due to the Y6W mutation results in changes up to 0.3 Å in the H-bond distances of several residues (N30, M92, R63 and D67) in the flavin environment (Figure 4i). The side chain of K29 also adopts a different orientation, pointing away from the flavin in the Y6W mutant compared to the WT OaPAC (Figure 4i). However, the most dramatic change is the ∼70° degree rotation around the CG – CD bond of the Q48 residue, with the distances of the amide nitrogen and carbonyl oxygen of the side chain of glutamine Q48 to the N_5_ of the flavin changing from 3.3 Å and 3.1 Å to 3.1 Å and 4.0 Å, respectively. The isoalloxazine moiety also rotates a few degrees in the Y6W mutant and displays an average shift of 0.3 Å, the side chain of M92 rotates ∼20°, whereas no notable changes are observed for W90 which retains a ‘W_out_’ orientation pointing away from the flavin as observed in the WT (Figure 4iii) and previously published OaPAC structures (Ohki *et al.,* 2016; Ohki *et al*., 2017; Chretien *et al*., 2024). The position of the rest of the nearby residues also does not change in the Y6W (Figure 4i). It should be pointed out that in order to explain the hydrogen bond rearrangement around the flavin in the WT protein upon blue-light excitation, a 180° rotation of the side chain of the glutamine residue and a keto-enol tautomerism of the glutamine residue both triggered by the electron and proton transfer from the tyrosine residue to the oxidized flavin have been proposed based on spectroscopic and computational studies for AppA and PixD (Tolentino Collado *et al*., 2024; Lukacs *et al*., 2022; Hontani *et al*., 2023; Hashem *et al*., 2023). In the recent TR-SFX study on the ATP-bound OaPAC a >180° rotation of the side chain Q48 was observed within a few microseconds after blue light excitation followed by a methionine/tryptophan switch (Chretien *et al*., 2024). The latter residues have received special attention as different conformations have been reported as discussed below and considered responsible for signal transduction in the BLUF domains. In particular, in AppA, the nearby methionine on the β5-strand of the BLUF domain has been shown to exist in two conformations, Met_in_ and Met_out_ (Jung *et al.,* 2006). Our studies on AppA_BLUF_ have revealed a dynamic picture for the tryptophan residue W104, with the dominant population pointing away from the flavin in the dark-adapted (17.7-20.5 Å) state and moving closer to the flavin in the light-adapted state (8.3-9.5 Å) (Karadi *et al*., 2020). On the contrary, recent solution ^19^F-NMR studies on AppA_BLUF_ have pointed towards a tryptophan (W104) entirely buried in the protein scaffold and a predominant W_in_ configuration in OaPAC_BLUF_ (Zhou *et al*., 2024). Recent MD simulations on the microsecond time-scale in AppA_BLUF_ have revealed an overall change in the structure of the BLUF domain with a sliding of the flavin ring within the pocket towards H85, a cleavage of the salt bridge of D82-R84 with subsequent formation of a new salt bridge between R84 and D28, an increase of the angle formed by the two α-helices and structural changes in the β-sheet originating from the glutamine Q63 and/or R84 (Hashem *et al*., 2023). In the AC domains of both WT and Y6W cryo-structures, there are two types of calcium binding sites (Figure S4, S5) as observed in our SFX OaPAC structure (Figure 2). At the internal calcium binding sites, calcium is coordinated with four (WT) and three (Y6W) water molecules and five amino acid residues (D156, D200, D270, N273, E279) (Figure S4iv/v, S5ii/iii). Calcium also binds at the surface of the protein forming bonds with three water molecules and two aminoacid residues (D310, D321) (Figure S4ii/iii, S5iv/v). Inspection of the AC domains in both cryo-structures reveals a 2.5 Å difference in the position of the calcium ions in the internal metal binding sites (Figure 5i). This difference suggests that a change in the H-bond environment of the flavin may affect the conformation of the AC domains and points out the allosteric regulation of the AC domains which can be triggered by a mutation (present case) or by light. The DDM map (Seno & Go, 1990) (Figure 5ii) also shows changes in the distances in the AC domain. In particular, in the Y6W OaPAC mutant, the BLUF domain is closer to the AC domain compared to the analogous distance in the wild-type OaPAC. Small differences are also observed in their internal cavities as illustrated in Figure 5iii, whereas the theoretical SAXS parameters do not show any differences between the two structures (Table 3).

**Figure 4.**
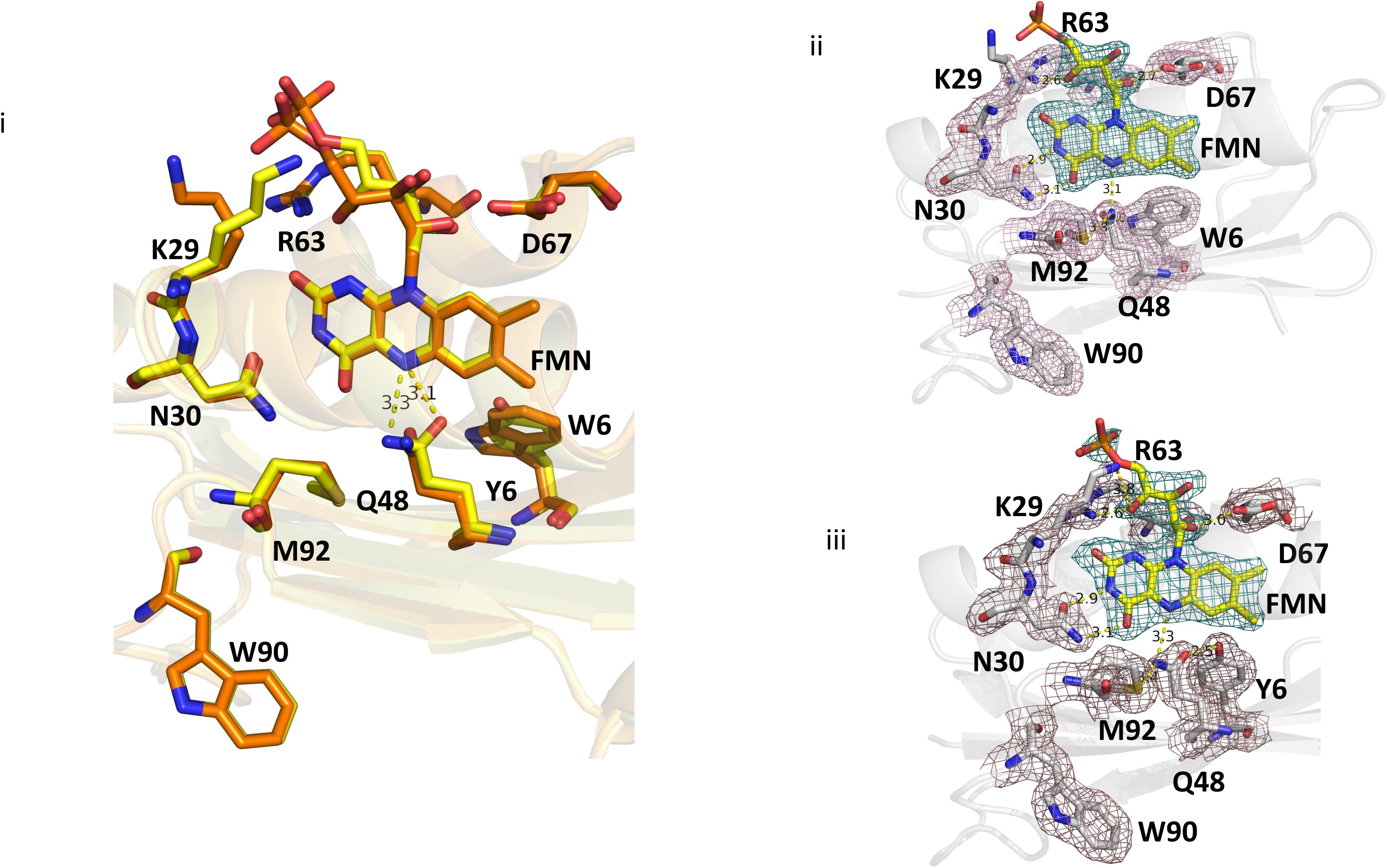
i) Superimposition of the BLUF domain of the cryo-WT (yellow) and Y6W OaPAC (orange). Distances are shown for the cryo-WT structure of OaPAC ii) Flavin active site of Y6W OaPAC in the cryo structure showing amino acid residues participating in the hydrogen bond network. 2Fo-Fc maps are represented in blue mesh (flavin) and purple (residues) at a contour level of 1.0 σ iii) Flavin active site of WT OaPAC in the cryo-structure showing amino acid residues participating in the hydrogen bond network. 2Fo-Fc maps are represented in blue mesh (flavin) and purple (residues) at a contour level of 1.0 σ.

**Figure 5.**
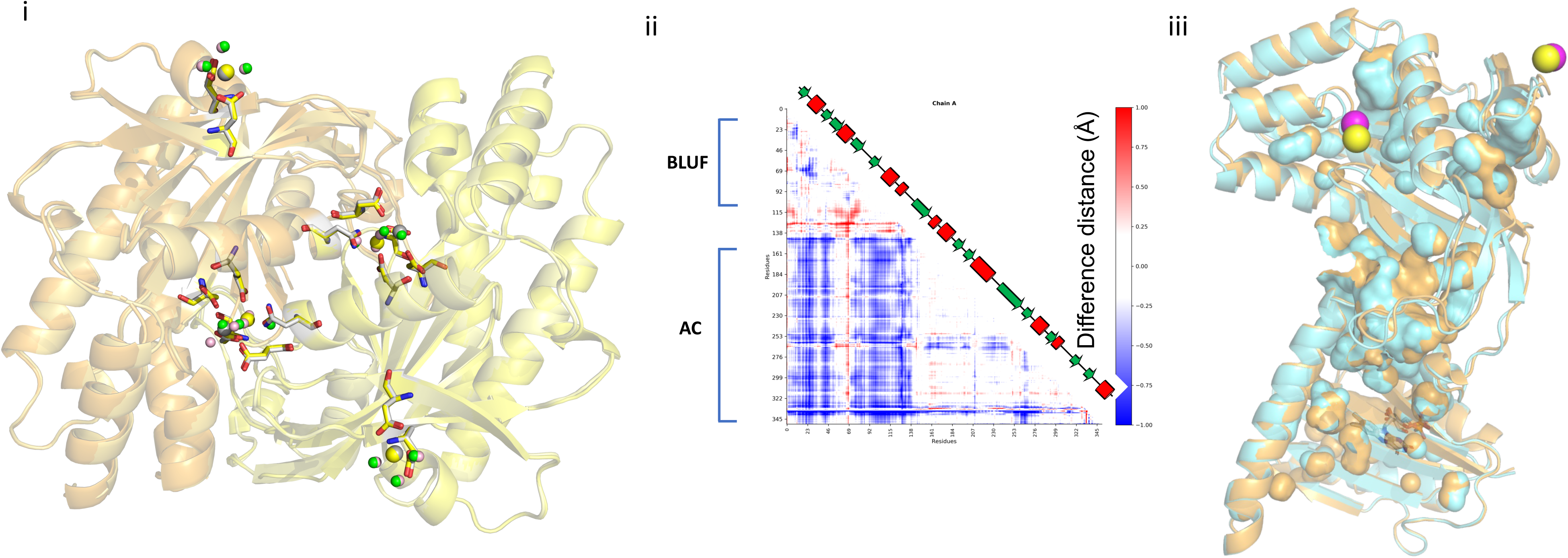
i) Superimposition of the AC domains of the cryo-structure of WT OaPAC and of the cryo-structure of Y6W OaPAC showing amino acid residues coordinated to the calcium ions (WT; grey spheres, Y6W; yellow spheres) and water molecules (WT; pink spheres, Y6W; green spheres) coordinated to calcium ions ii) DDM map calculated between the cryo-structure of WT OaPAC and Y6W OapAC mutant. Red and blue indicate increasing and decreasing distance, respectively with respect to the cryo-structure of wt OaPAC iii) Superimposition of monomeric OaPAC (WT orange; Y6W; cyan; calcium ions in WT; pink spheres, calcium ions in Y6W; yellow) showing cavities calculated with PyMOL.

### 3.6 Effect of the crystallization condition on the crystal structure of OaPAC

Comparison of our OaPAC structures with the recently released OaPAC structures of Chretien *et al*. (Chretien *et al*., 2024) and the available OaPAC structures of Ohki *et al*. (Ohki *et al*. 2016; 2017) reveals some interesting features, presumably arising from the different crystallization conditions used (this work; 0.06 M divalents, 0.1 M Tris-Bicine pH 8.5, 30% v/v PEG550 MME-PEG20000; Chretien *et al*.; 100 mM SPG buffer pH 7.0, 1.2 M disodium succinate, 100 mM guanidine HCl and 5 mM MgCl_2_, Ohki *et al*.; 0.1 M sodium citrate pH 5.0, 10% PEG 20K, 5 mM magnesium chloride, 5 mM ApCpp). As discussed in section 3.1, OaPAC has been crystallized in various crystal forms: orthorhombic C222_1_, pdb:8qfe, 8qff, 8qfh; orthorhombic P2_1_2_1_2_1_, pdb:4yut; and hexagonal P6_1_22, pdb:4yus. In our preparation, OaPAC has crystallized in the orthorhombic space group P2_1_2_1_2. Figure 6 which compares the AC domains of our cryo-OaPAC structure with the SFX ATP bound OaPAC structure of Chretien *et al*. reveals a significant reorientation of a helix in the ATP binding domain in our conditions (Figure 6i). This reorientation seems to originate from a loop placed on top of the α5 helix which in our preparation seems to push down the α5 helix resulting in the observed kink of the helix. As a result, the accessible solvent area (ASA) for the ATP pocket, as calculated using the PISA server (https://www.ebi.ac.uk/pdbe/pisa/) and a custom-built python script, is reduced from 511 Å^2^ in the cryo-structure of Chretien *et al*. (pdb: 8qfe) (Figure S6i) to 315 Å^2^ in our cryo-OaPAC structure (Figure 6iii/S6iv). This is a significant reduction which results in a close conformation that may prevent ATP from binding. Notably, ATP can bind to OaPAC crystallized in conditions supporting an open conformation (RT ATP-bound OaPAC; pdb: 8qfh) (Figure 6ii/S6v). Similar to the structures of Chretien *et al*. (Chretien *et al*. 2024) those reported by Ohki *et al*. (Ohki *et al*. 2016; 2017) also display an open conformation for the ATP pocket (Figure S6ii and S6iii) compared to our cryo-OaPAC structure. A B-factor putty representation showing differences in the B-factors between the available OaPAC structures is shown in Figure S7. These differences reflect the effect that ligand binding and various crystallization conditions and temperatures can have in the structure of OaPAC.

**Figure 6.**
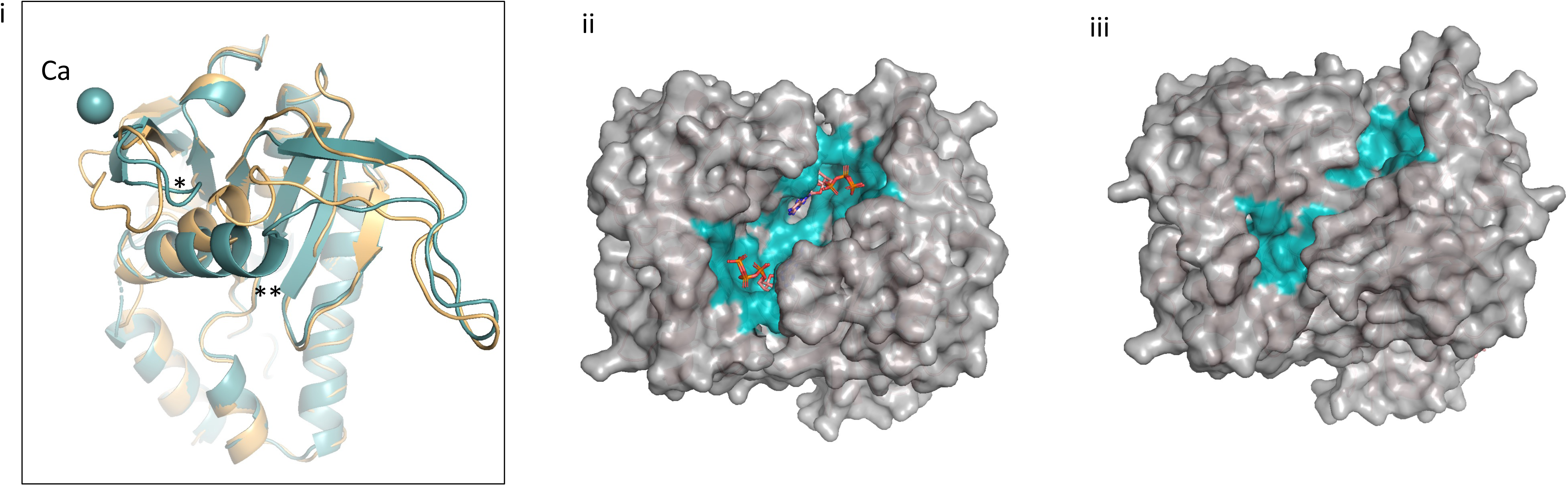
i) Comparison of our cryo-OaPAC structure (green) with the ATP-bound SFX structure (pdb: 8qfh, yellow orange) showing the loop (labelled with an asterisk) that pushes down a helix (labelled with two asterisks) ii) ATP binding pocket in the ATP-bound OaPAC structure of Chretien *et al*. (pdb (8qfh) iii) ATP binding pocket in our cryo-OaPAC structure.

## Conclusions

This study paves the way for time-resolved studies at XFELs (TR-SFX) and synchrotrons (TR-SSX) which will provide structural information on the early intermediates (ps-ms) forming upon blue-light excitation of the substrate- and radiation damage-free OaPAC. These structures will further allow a correlation of the available spectroscopic signatures on the substrate-free enzyme (Collado *et al*., 2022, Tolentino Collado *et al.,* 2024) with the early-formed structural intermediates, making it possible to obtain a complete picture of the photoactivation mechanism of the enzyme. This study also highlights how the crystallization conditions can influence crystal packing which may result in conformational differences affecting ligand binding and how changes in the sensor domain of a protein can affect allosterically its effector domain.

## Acknowledgements

This project has received funding from the European Union’s Horizon 2020 research and innovation programme under the Marie Skłodowska-Curie grant agreement No 893817 (IF to S.M.K). S.M.K acknowledges Peter C. E. Moody for an inspiring seminar on XFELs, Sylvain Engilberge for discussions on protein crystallization and Sofia Jaho and Kyprianos Hadjidemetriou for critical reading of the manuscript. This work was partially carried out at the platforms of the Grenoble Instruct-ERIC center (IBS and ISBG; UMS 3518 CNRS-CEA-UGA-EMBL) within the Grenoble Partnership for Structural Biology (PSB). Platform access was supported by FRISBI (ANR-10-INBS-05-02) and GRAL, a project of the University Grenoble Alpes graduate school (Écoles Universitaires de Recherche) CBH-EUR-GS (ANR-17-EURE-0003). The IBS acknowledges integration into the Interdisciplinary Research Institute of Grenoble (IRIG, CEA). We acknowledge European XFEL in Schenefeld, Germany for provision of X-ray free-electron laser beamtime at SPB/SFX SASE1 under proposal number 3018 (PCS beamtime) and we would like to thank the staff for their assistance. Data recorded for the experiment at the European XFEL are available at doi: 10.22003/XFEL.EU-DATA-003018-00. S.M.K. acknowledges IBS BAGs and the PIXE beamtime at the Surrey Ion Beam Centre under the RADIATE program (proposal 23003164-ST).

---

**Figure S1.** UV-vis spectra of OaPAC in the dark-(black line) and light-adapted state (red line) in the crystalline form (i) and in solution (ii).

**Figure S2.** Ion modelling in OaPAC crystal structures.

Based on crystallisation conditions, either magnesium or calcium ions could be modelled into specific region of the protein. Two models were generated (one with each type of ions) and refined using the EuXFEL RT diffraction data. Resulting signal in the mFo-DFc maps is displayed as green (contoured at +3.5 sigma) and red (contoured at 3.5 sigma) meshes. Panels i) and ii) represent the modelling of Mg^2+^ ion (orange spheres) in the two different regions. The protein model is represented as teal sticks, and water molecules as red spheres. The 2mFo-DFc is represented as an isosurface contoured at 1.5 sigma. The green density is indicative of a lack of electrons. Panels iii) and iv) represent models with calcium ions (grey sphere), without any sign of signal in the mFo-DFc map. Such comparison favours the modelling of calcium ions in the structures.

**Figure S3.** Theoretical SAXS data derived from the available crystal structures of OaPAC using the FoXS server. A) SFX, B) SSX, C) cryo-, D) Y6W. i) theoretical SAXS profile ii) Guinier analysis iii) Kratky plot iv) Pair distribution function, P(r).

**Figure S4.** i) AC domain of cryo-wt OaPAC showing amino acid residues coordinated to the calcium ions (green spheres). Water molecules coordinated to the calcium ions are shown as red spheres. ii) and iii) Zoom-in on the calcium binding sites on the surface of wt OaPAC iv) and v) Zoom-in on the dimer interface and the calcium binding sites of wt OaPAC; the 2Fo-Fc map is coloured in blue and shown only for the calcium ions.

**Figure S5.** i) AC domain of cryo-Y6W OaPAC showing amino acid residues coordinated to the calcium ions (green spheres). Water molecules coordinated to the calcium ions are shown as yellow spheres. ii) and iii) Zoom-in on the dimer interface and the calcium binding sites of Y6W OaPAC. iv) and v) Zoom-in on the calcium binding sites on the surface of Y6W OaPAC iv) and v) the 2Fo-Fc map is colored in blue and shown only for the calcium ions (green spheres; water molecules: red spheres).

**Figure S6.** Comparison of the ATP binding pocket shown for the cryo-ATP-free OaPAC structure of Chretien *et al*. (pdb: 8qfe) (511 Å), ii) OaPAC pdb: 4yut, iii) OaPAC pdb: 4yus, iv) our cryo-OaPAC structure pdb: 9f1y (315 Å), v) SFX ATP-bound OaPAC structure of Chretien *et al*. (pdb: 8qfh), vi) SFX OaPAC structure pdb: 9f1w (378 Å), vii) SSX OaPAC structure pdb: 9f1x (263 Å) and viii) cryo-OaPAC Y6W structure pdb: 9f1z (282 Å). In parenthesis is given, the accessible solvent area (ASA) for the ATP pocket.

**Figure S7.** Comparison of the B-factor putty representations for all OaPAC structures available.

**Appendix.** Video showing jetting of the slurry of OaPAC microcrystals tested at the XBI lab (EuXFEL).

## Notes

https://youtu.be/KZbVdYs0lLs

### Competing Interest Statement

The authors have declared no competing interest.

