## Supplemental Figures for "Crystal structure of a bacterial photoactivated adenylate cyclase determined at room temperature by serial femtosecond crystallography"

[c] European XFEL, Holzkoppel 4, 22869 Schenefeld, Germany.

[d] Department of Biophysics, Medical School, University of Pecs, Szigeti Street 12, Pécs 7624 Hungary

[e] University of Surrey Ion Beam Centre, Guildford, GU2 7XH United Kingdom.

[f] Institute for Nanostructure and Solid-State Physics, Universität Hamburg, HARBOR, Luruper Chaussee 149, 22761 Hamburg, Germany.

[g] Department of Biochemistry, University of Oxford, Dorothy Crowfoot Hodgkin Building, South Parks Road, Oxford OX1 3QU, United Kingdom.

### Current address: Department of Biophysics, Medical School, University of Pécs, Szigeti Street 12, Pecs 7624, Hungary

§ Current address: Center for Free-Electron Laser Science CFEL, Deutsches Elektronen-Synchrotron DESY, Notkestr. 85, 22607

Hamburg, Germany

Figure S1

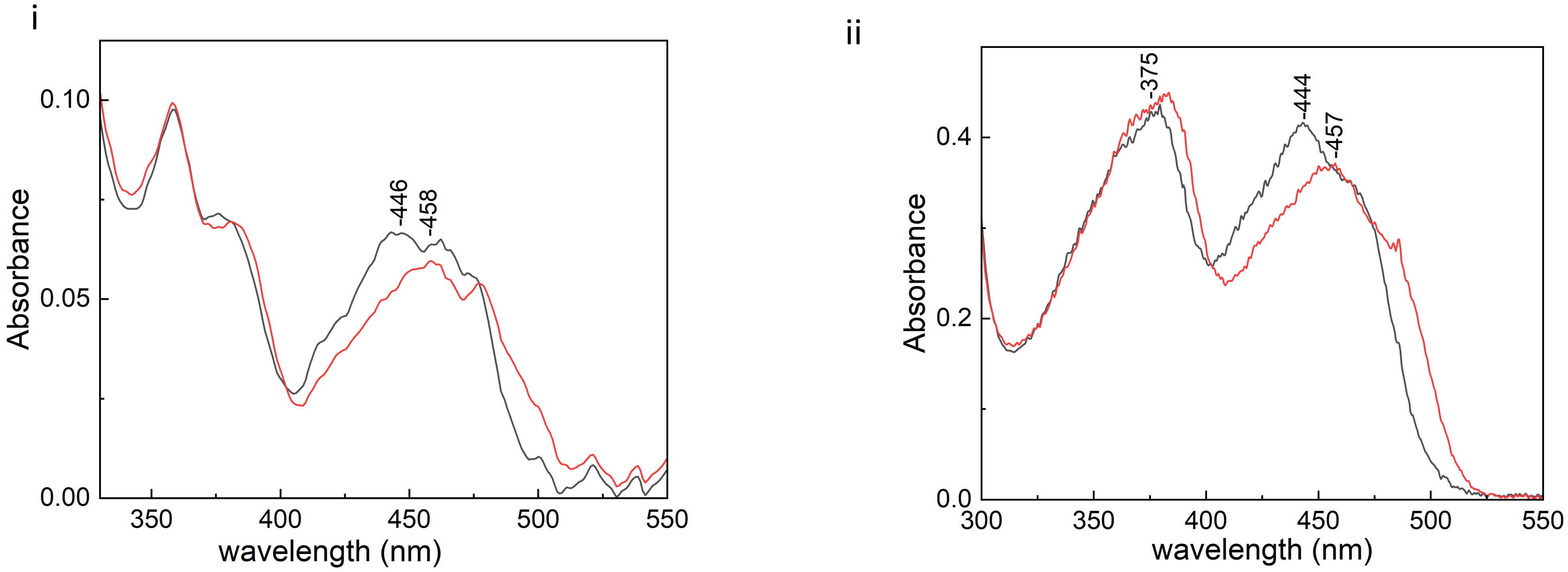
i
0.10
0.05
0.00
Absorbance
350
400
450
500
550
wavelength (nm)
-446
-458
ii
0.4
0.2
0.0
Absorbance
300
350
400
450
500
550
wavelength (nm)
-375
-444
-457

Figure S2

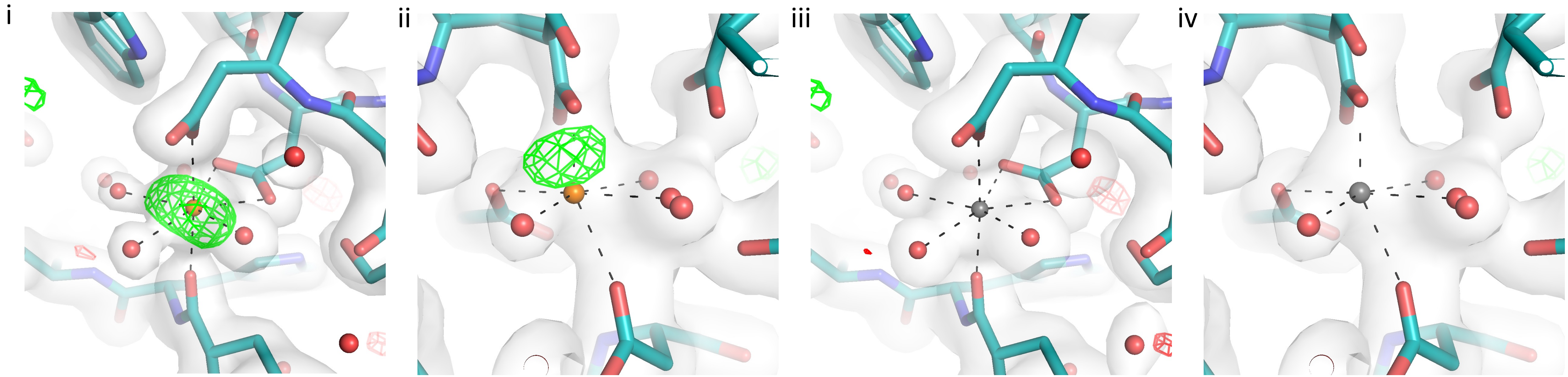

Figure S3

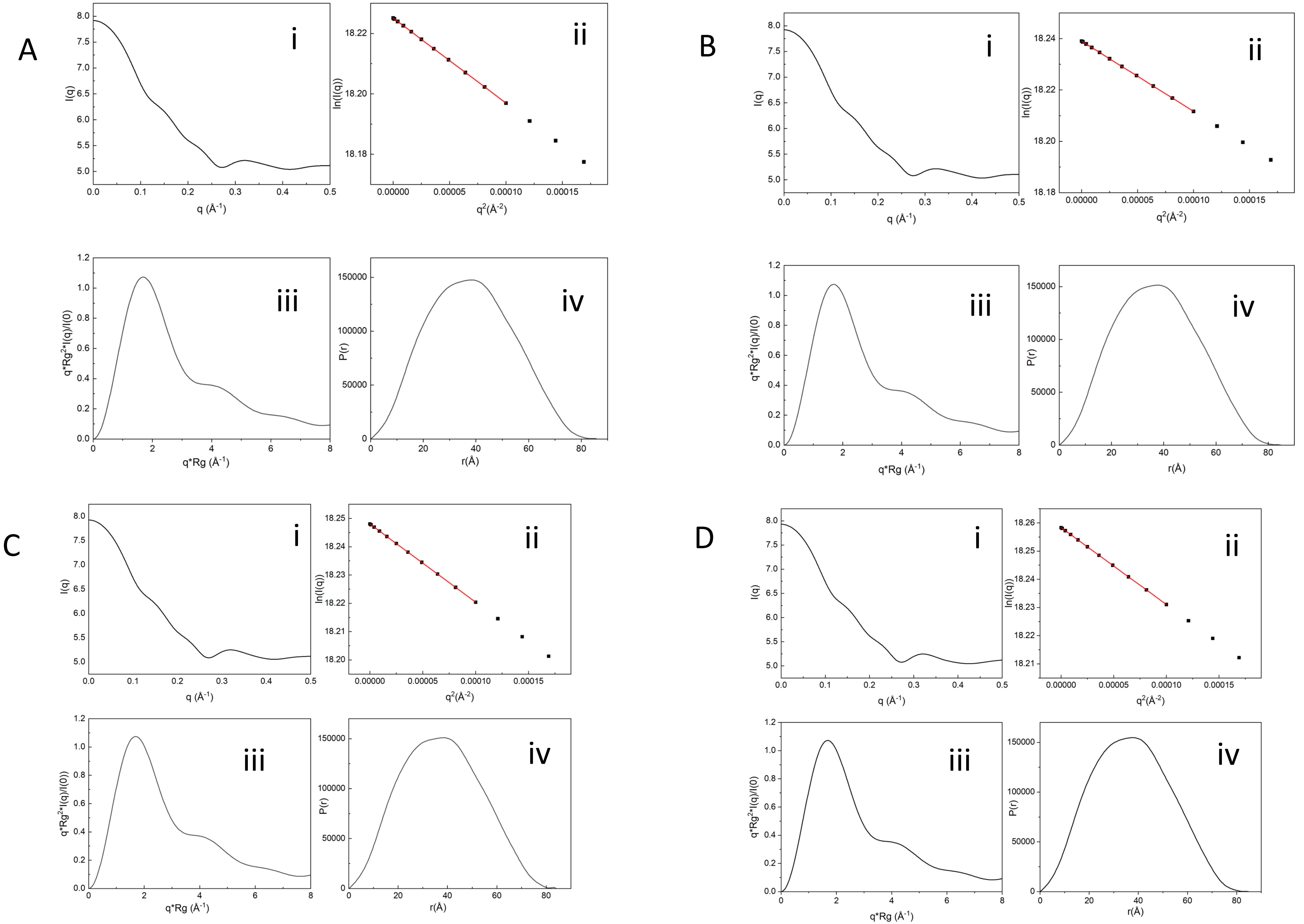

Figure S4

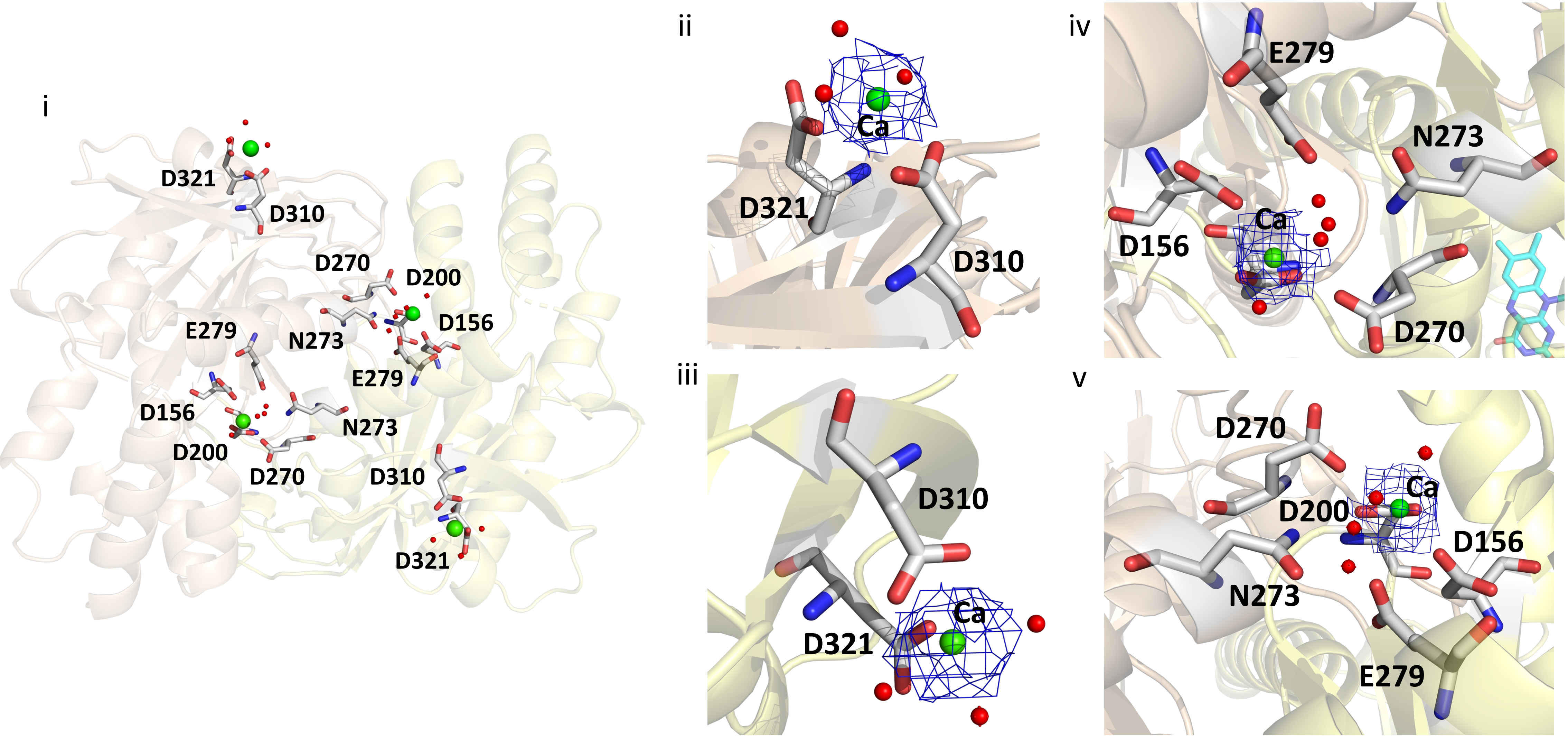
i
D321
D310
D270
D200
D156
E279
N273
E279
D156
N273
D200
D270
D310
D321
ii
Ca
D321
D310
iii
D310
D321
Ca
iv
E279
N273
Ca
D156
D270
v
D270
Ca
D200
D156
N273
E279

Figure S5

i

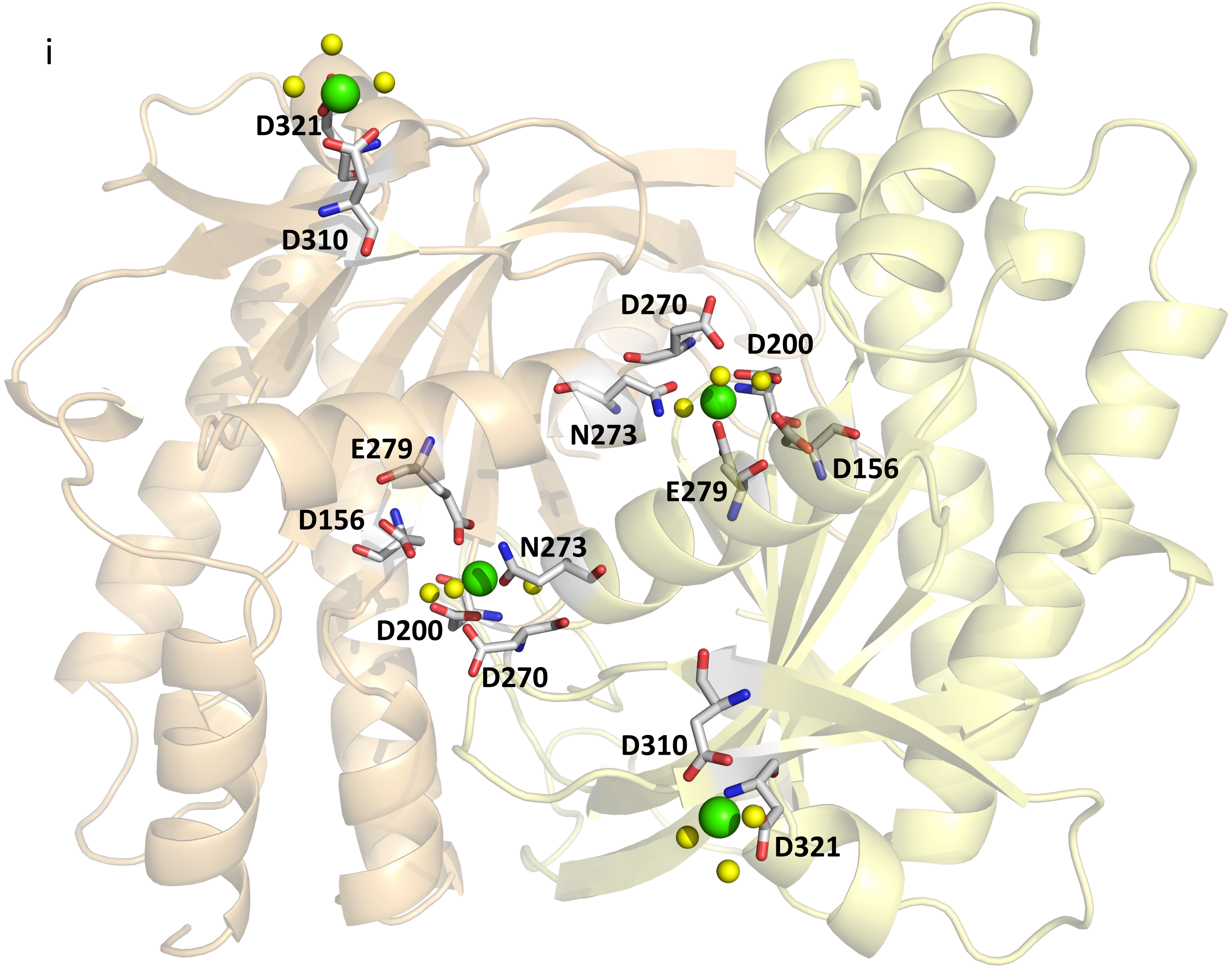
D321
D310
D270
D200
N273
E279
D156
E279
D156
N273
D200
D270
D310
D321

Figure S5

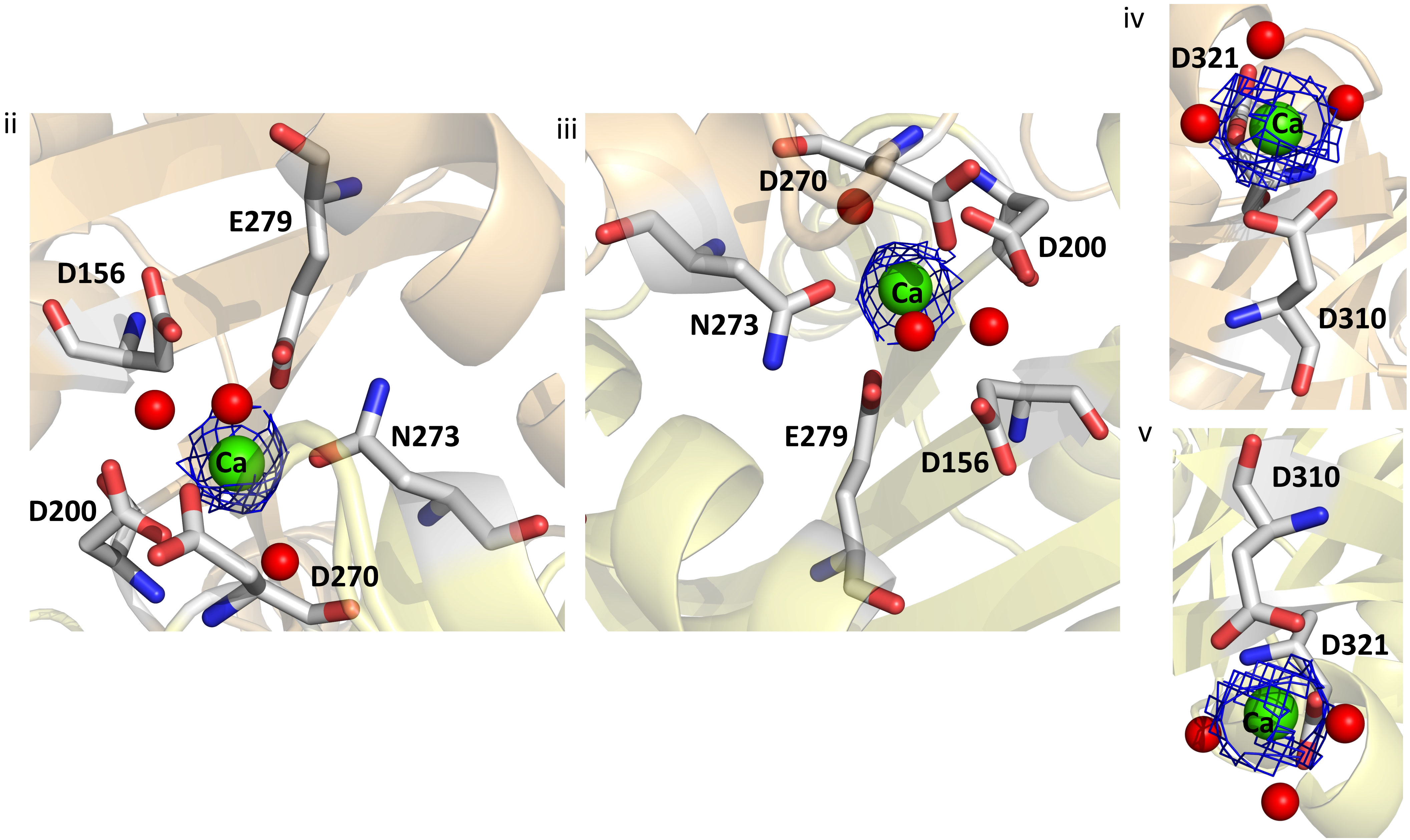

ii
E279
D156
Ca
N273
D200
D270
iii
D270
D200
N273
Ca
E279
D156
iv
D321
Ca
D310
v
D310
D321
Ca

Figure S6

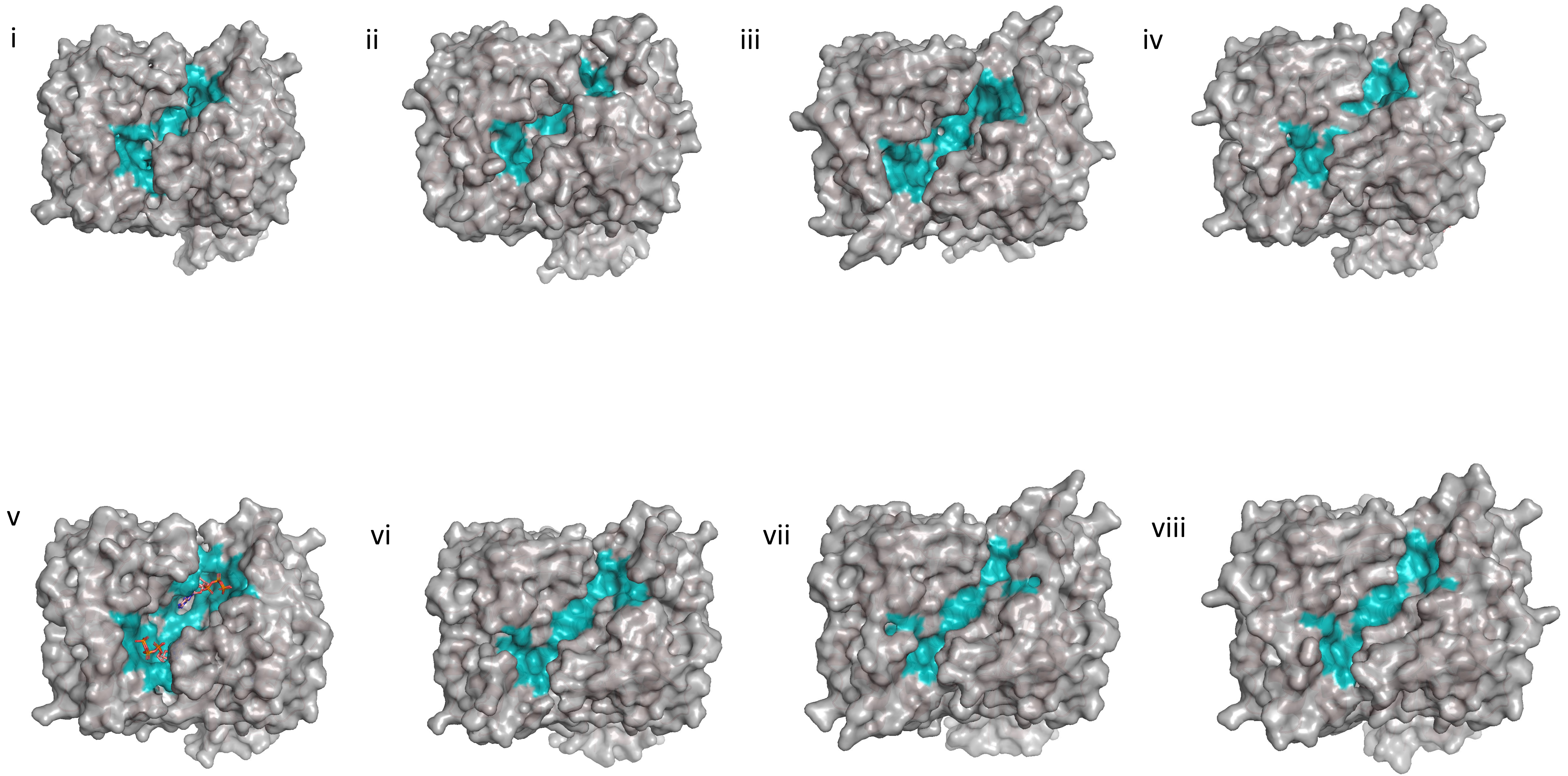

Figure S7

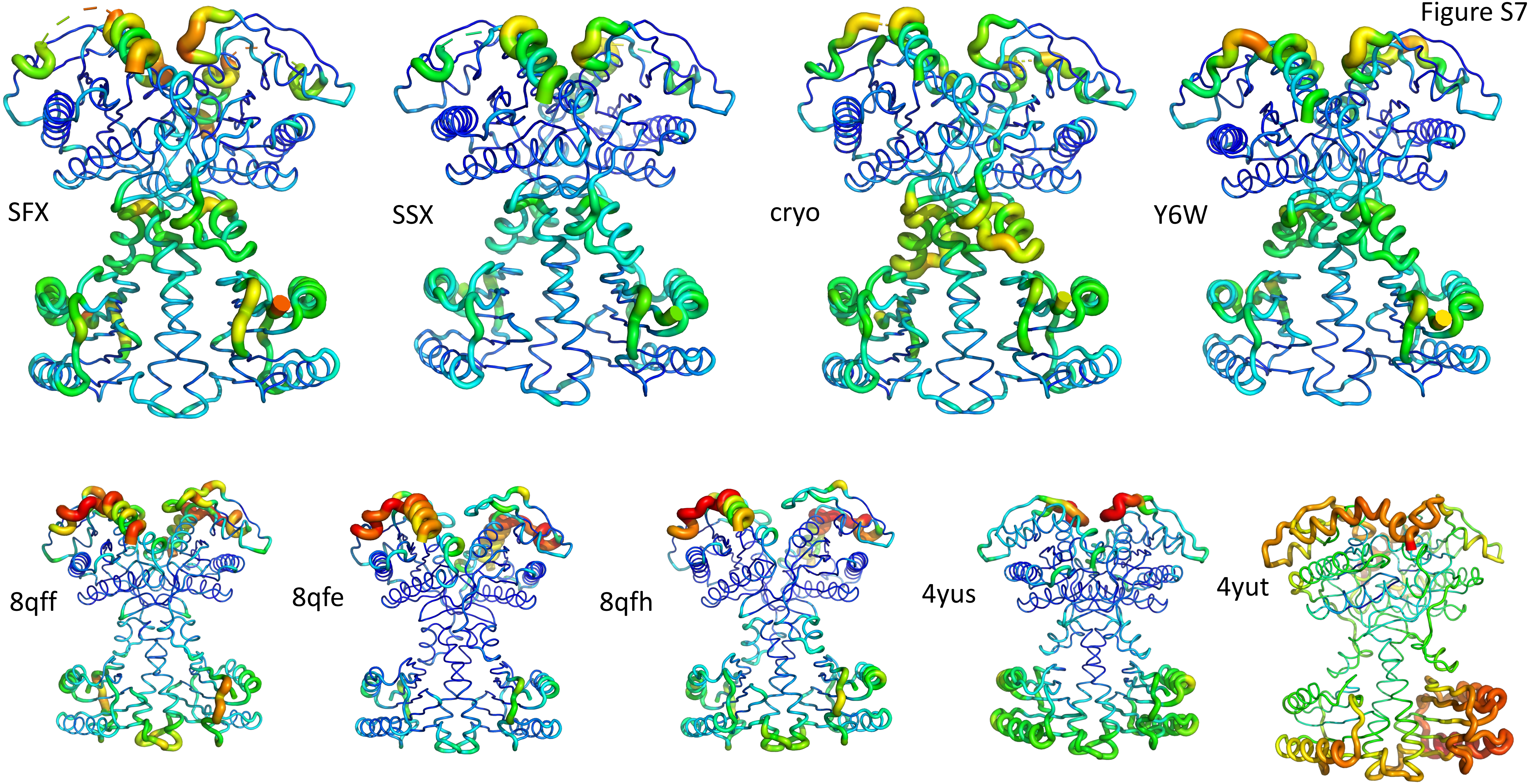
